# Expansion of Disease Specific Cardiac Macrophages in Immune Checkpoint Inhibitor Myocarditis

**DOI:** 10.1101/2023.04.28.538426

**Authors:** Pan Ma, Jing Liu, Juan Qin, Lulu Lai, Gyu Seong Heo, Hannah Luehmann, Deborah Sultan, Andrea Bredemeyer, Geetika Bajapa, Guoshuai Feng, Jesus Jimenez, Antanisha Parks, Junedh Amrute, Ana Villanueva, Yongjian Liu, Chieh-Yu Lin, Matthias Mack, Kaushik Amancherla, Javid Moslehi, Kory J. Lavine

## Abstract

**Background:** Immune checkpoint inhibitors (ICIs), antibodies targeting PD-1/PD-L1 or CTLA4 have revolutionized cancer management but are associated with devastating immune-related adverse events (irAEs) including myocarditis. The main risk factor for ICI myocarditis is the use of combination PD-1 and CTLA4 inhibition. ICI-myocarditis is often fulminant and is pathologically characterized by myocardial infiltration of T lymphocytes and macrophages. While much has been learned regarding the role of T-cells in ICI-myocarditis, little is understood regarding the identity, transcriptional diversity, and functions of infiltrating macrophages.

**Methods:** We employed an established murine ICI myocarditis model (*Ctla4^+/-^Pdcd1^-/-^* mice) to explore the cardiac immune landscape using single-cell RNA-sequencing, immunostaining, flow cytometry, in situ RNA hybridization and molecular imaging and antibody neutralization studies.

**Results:** We observed marked increases in CCR2^+^ monocyte-derived macrophages and CD8^+^ T-cells in this model. The macrophage compartment was heterogeneous and displayed marked enrichment in an inflammatory CCR2^+^ subpopulation highly expressing *Cxcl9*, *Cxcl10*, *Gbp2b*, and *Fcgr4* that originated from CCR2^+^ monocytes. Importantly, a similar macrophage population expressing *CXCL9*, *CXCL10*, and CD16α (human homologue of mouse FcgR4) was found selectively expanded in patients with ICI myocarditis compared to other forms of heart failure and myocarditis. *In silico* prediction of cell-cell communication suggested interactions between T-cells and *Cxcl9^+^Cxcl10^+^* macrophages via IFN-γ and CXCR3 signaling pathways. Depleting CD8^+^T-cells, macrophages, and blockade of IFN-γ signaling blunted the expansion of *Cxcl9^+^Cxcl10^+^*macrophages in the heart and attenuated myocarditis suggesting that this interaction was necessary for disease pathogenesis.

**Conclusion:** These data demonstrate that ICI-myocarditis is associated with the expansion of a specific population of IFN-γ induced inflammatory macrophages and suggest the possibility that IFN-γ blockade may be considered as a treatment option for this devastating condition.

## Introduction

Immune checkpoint inhibitors (ICIs) have revolutionized cancer therapy. Remarkable improvements in clinical response rates, tumor-free, and overall survival have led to the widespread use of these agents across various malignancies^1–3^. ICIs are now combined where two separate immune checkpoints e.g., PD-1/PD-L1 and CTLA4 or LAG3 are inhibited with demonstration of greater anti-tumor efficacy. However, ICIs especially when used in combination are associated with a wide spectrum of immune-related adverse events (irAEs) which can affect any organ^4, 5^. While infrequent, myocarditis is the most serious irAE with high mortality despite corticosteroid therapy. ICI myocarditis is characterized by T-cell and macrophage infiltration with associated cardiomyocyte death. Patients often present with electrocardiographic disturbances including conduction block and ventricular arrhythmias; 50% of patients have a normal systolic cardiac function^6, 7^. Combined use of PD-1 and CTLA4 inhibitors is the major risk factor for myocarditis. Higher frequency of myocarditis was reported in patients treated with combination treatment (1.22%) compared with PD-1/PD-L1 or CTLA4 inhibitors alone (0.54%)^8^. Previous studies addressing ICI myocarditis pathogenesis primarily focused on T-cells^9–12^. However, less is known about macrophage populations in ICI myocarditis and potential communication between immune cell types, such as T-cells and macrophages.

Under steady state conditions, the heart contains predominantly macrophages with smaller populations of T-cells, B-cells, and dendritic cells^13^. Cardiac macrophages represent a heterogeneous population of cells that can be divided into two major subsets with differing origins and functions: cardiac resident CCR2^-^ macrophages and monocyte-derived CCR2^+^ macrophages^14–16^. Among these subsets, CCR2^+^ macrophages display the greatest inflammatory potential and contribute to myocardial inflammation and heart failure pathogenesis. CCR2^+^ macrophages generate numerous inflammatory mediators (cytokines, chemokines, oxidative products) and function as antigen presenting cells. In experimental acute autoimmune myocarditis, CCR2 silencing attenuates disease^17, 18^.

It is well recognized that activated T-cells communicate with macrophages as a component of the host immune response in the setting of infection and autoimmunity^19^. Nevertheless, how macrophages contribute to ICI myocarditis and to what degree macrophage-T cell crosstalk modulate this process is not clear. Immune checkpoint therapies including PD-1 and CTLA4 inhibitors boost T-cell responses and promote abundant cytokine production^20^. Within the context of ICI myocarditis, interactions between activated T-cells and cardiac macrophages may drive myocardial inflammation and contribute to disease pathogenesis. This possibility remains to be explored.

Here, we dissected the cardiac immune landscape of ICI myocarditis using an established mouse model (*Ctla4^+/-^Pdcd1^-/-^*mice) by employing immunostaining, flow cytometry, in situ RNA hybridization, single-cell RNA-sequencing, molecular imaging and antibody neutralization studies. We observed evidence of CD8^+^ T-cell activation and expansion of an inflammatory CCR2^+^ monocyte-derived macrophage population that displayed a transcriptional signature characterized by *Cxcl9*, *Cxcl10*, *Gbp2b*, and *Fcgr4* expression that originated from CCR2^+^ monocytes. A similar population of macrophages expressing *CXCL9*, *CXCL10*, and CD16α (human homologue of mouse FcgR4) were found selectively expanded in patients with ICI myocarditis compared to other forms of heart failure and myocarditis. Informatic prediction of cell-cell communication suggested that T-cells regulate the expansion of *Cxcl9^+^Cxcl10^+^* macrophages via IFN-γ signaling and highlighted a positive feedback loop between these cell types mediated by *CXCL9*, *CXCL10*, and CXCR3 signaling. Consistent with this prediction, depleting CD8^+^T-cells, macrophages, and blockade of IFN-γ signaling blunted the expansion of *Cxcl9^+^Cxcl10^+^* macrophages in the heart and prolonged survival. Collectively, these data demonstrate that ICI myocarditis is associated with the expansion of a specific population of IFN-γ induced inflammatory macrophages and suggest the possibility that IFN-γ blockade may serve as a potential intervention strategy for this devastating condition.

## Materials and Methods

### Mouse strains

All animal studies were performed in compliance with guidelines set forth by the National Institutes of Health Office of Laboratory Animal Welfare and approved by the Washington University institutional animal care and use committee. All mouse strains utilized (wild type, *Ctla4^+/+^Pdcd1^-/-^*, and *Ctla4^+/-^Pdcd1^-/-^* mice) were on the C57BL/6 background. All mice were genotyped according to established protocols. Female mice were primarily employed in the experiments.

### Study approval and human pathological specimens

This study was approved by the Institutional Review Board of Washington University in St. Louis. Hematoxylin and eosin-stained slides were reviewed by cardiovascular pathologists to confirm the diagnosis.

### Statistical analyses

Normal distribution of quantification results was tested by Shapiro-Wilk test. Significance of the quantification results was tested by Mann-Whitney test or Welch’s t-test or Unpaired t-test using Prism 9.0 (GraphPad Software, San Diego, CA) or using R package Seurat, ClusterProfiler, Cellchat. P-value <0.05 was considered as significant.

### Data and codes availability

All the codes used in the manuscript are deposited in GitHub (<u>pma22wustl/Expansion-of-Disease-Specific-Cardiac-Macrophages-in-Immune-Checkpoint-Inhibitor-Myocarditis (github.com)</u>). Other data available upon reasonable request from the Lead Contact. The single-cell raw expression matrices and raw sequence files that support the findings of this study are available on the Gene Expression Omnibus (GSE 227437, GSE230192). Published RNA-seq data of male mice that support the findings of this study are available on the Gene Expression Omnibus (GSE225099). Published RNA-seq data of ICI associated myocarditis patients are available in the EBI ArrayExpress Database, accession number E-MTAB-8867^21^.

## Results

### Accumulation of CCR2^+^ macrophages in a mouse model of ICI myocarditis

To dissect the cardiac immune landscape of ICI myocarditis, we employed a previously validated mouse model of this condition whereby haploinsufficiency in *Ctla4* in the background of PD-1 deficiency (*Pdcd1^-/-^*) results in sudden cardiac death and pathological characteristics consistent with ICI myocarditis. Clinical features include electrocardiographic disturbances with relatively preserved cardiac function consistent with human disease. Immune infiltration is limited to a few tissues including the heart, with higher prevalence of disease in female mice ^8^. As such, given the variability of presentation with myocarditis, female mice were primarily used in this study. Consistent with previous reports^12^, we detected robust immune cell accumulation in the hearts of *Ctla4^+/-^Pdcd1^-/-^* mice with increased abundance of T-cells (CD8^+^ > CD4^+^) (**Supplementary Fig 1A**) and CD68^+^ macrophages (**Fig 1A**) compared with wild type mice. *Ctla4^+/+^Pdcd1^-/-^* mice displayed an intermediate phenotype. Flow cytometry demonstrated increased macrophage, CD8^+^ T-cell and CD4^+^ T-cell abundance in *Ctla4^+/-^Pdcd1^-/-^* hearts compared to *Ctla4^+/+^Pdcd1^-/-^* hearts (**Fig 1B)**. NK-cells and B-cells were not affected **(Supplementary Fig 1B**). Further phenotyping of the monocyte/macrophage compartment revealed increased frequency of LY-6C^high^ monocytes and CCR2^+^ macrophages in *Ctla4^+/-^Pdcd1^-/-^* hearts compared to *Ctla4^+/+^Pdcd1^-/-^* hearts (**Fig 1C**). Using *in situ* hybridization, we detected robust *Ccr2* mRNA expression in *Cd68*^+^ macrophages in *Ctla4^+/-^Pdcd1^-/-^* hearts (**Fig 1D**). These findings were confirmed by CCR2 targeted positron emission tomography/computed tomography (PET/CT) imaging as previously demonstrated^22^. ^68^Ga-DOTA-ECL1i PET/CT revealed increased uptake in the hearts of *Ctla4^+/-^Pdcd1^-/-^ mice* compared to wild type and *Ctla4^+/+^Pdcd1^-/-^* hearts (**Fig 1E**). Collectively, these findings reveal that ICI myocarditis is associated with the accumulation of CCR2^+^ macrophages.

**Fig 1.**
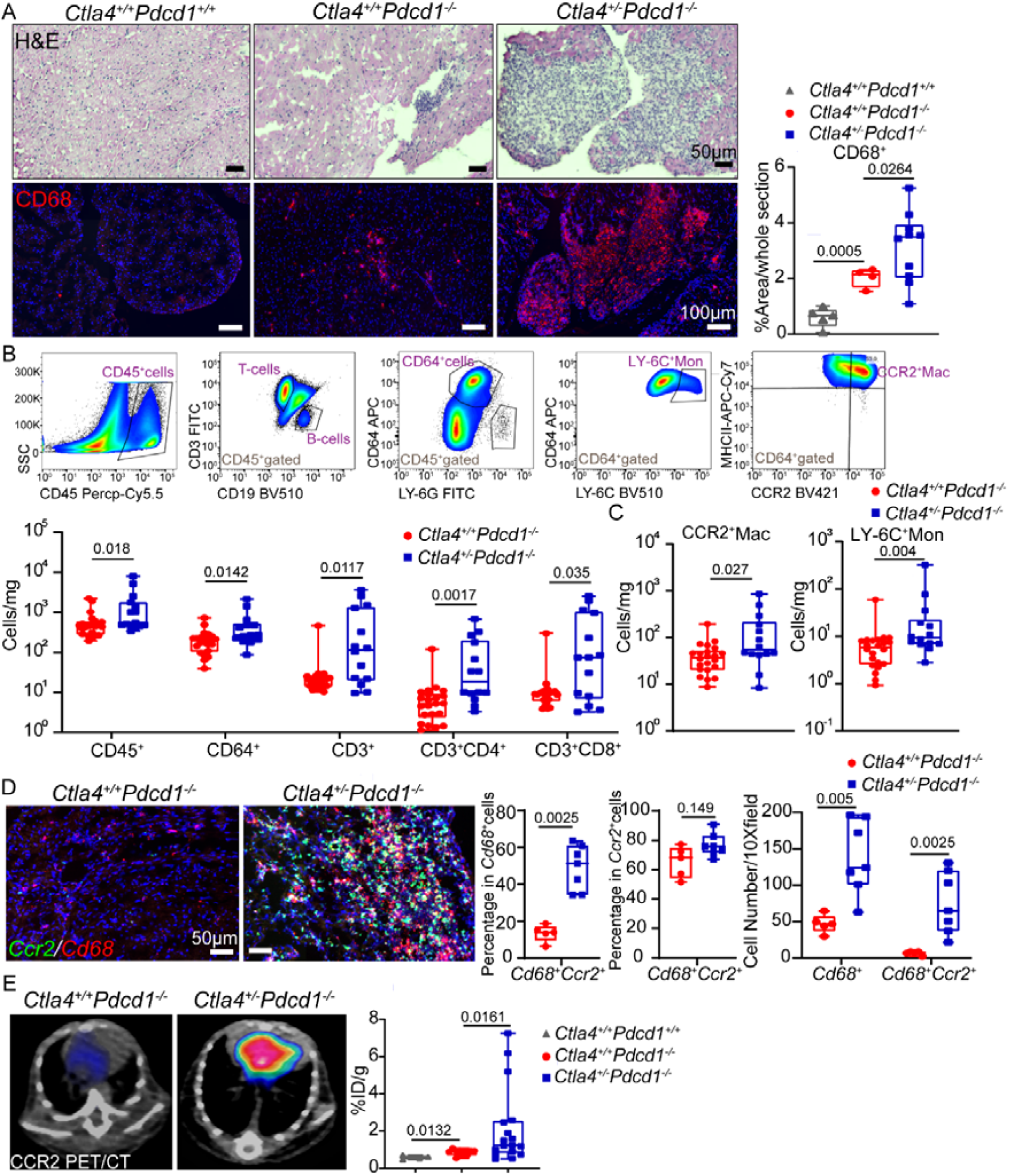
Accumulation of CCR2^+^ monocytes and macrophages in *Ctla4^+/-^Pdcd1^-/-^*mouse hearts. (A) Representative images of H&E and CD68 immunofluorescent staining (red) in wild type (*Ctla4^+/+^Pdcd1^+/+^*), *Ctla4^+/+^Pdcd1^-/-^*, and *Ctla4^+/-^Pdcd1^-/-^*hearts. Quantification of CD68^+^ cells. Data collected from two independent experiments. *Ctla4^+/+^Pdcd1^+/+^* (n=5), *Ctla4^+/+^Pdcd1^-/-^*(n=4), *Ctla4^+/-^Pdcd1^-/-^* (n=10), Welch’s t test, two-tailed. Scale bar for H&E staining images, 50 µm. Scale bar for CD68 staining images, 100 µm (B) Quantification of CD45^+^, CD64^+^, CD3^+^, CD3^+^CD4^+^, CD3^+^CD8^+^ cells in the heart by flow cytometry. Data collected from four independent experiments. *Ctla4^+/+^Pdcd1^-/-^* (n=22), *Ctla4^+/-^Pdcd1^-/-^*(n=14), Mann-Whitney test, two-tailed. (C) Quantification of CCR2^+^ macrophages and LY-6C^high^ monocytes by flow cytometry. Data collected from four independent experiments. *Ctla4^+/+^Pdcd1^-/-^*(n=22), *Ctla4^+/-^Pdcd1^-/-^* (n=14), Mann-Whitney test, two-tailed. (D) *Ccr2* (green) and *Cd68* (red) expression detected in *Ctla4^+/+^Pdcd1^-/-^* and *Ctla4^+/-^ Pdcd1^-/-^* mouse hearts via RNA in situ hybridization. (Left) Representative images, scale bar, 50 µm. (Right) Quantification of the percentage of *Cd68^+^Ccr2^+^*cells in *Cd68^+^*cells and *Ccr2^+^*cells, respectively as well as the cell numbers per 10X field. *Ctla4^+/+^Pdcd1^-/-^* (n=5), *Ctla4^+/-^Pdcd1^-/-^* (n=7), Mann-Whitney test, two-tailed. (E) *In vivo* cardiac CCR2 signal was detected with a CCR2 specific radiotracer, ^64^Cu-DOTA-ECL1i using positron emission tomography (PET). Representative CCR2 PET/CT images (left) and quantification of CCR2 tracer uptake (right). Data collected from two independent experiments, *Ctla4^+/+^Pdcd1^+/+^* (n=4), *Ctla4^+/+^Pdcd1^-/-^* (n=12), *Ctla4^+/-^Pdcd1^-/-^* (n=17), Mann-Whitney test, two-tailed.

### Expansion of a unique population of CCR2^+^ macrophages in a mouse model of ICI myocarditis

To explore the cardiac immune landscape of ICI myocarditis, we performed single-cell RNA-sequencing (scRNA-seq) of heart tissue from control (*Ctla4^+/+^Pdcd1^-/-^*, n=4) and ICI myocarditis (*Ctla4^+/-^Pdcd1^-/-^*, n=10) mice. We constructed five libraries (control: L1, L2; disease: L3, L4, L5) from DR^+^DAPI^-^ cells purified from pooled enzymatically digested hearts by FACS (**Supplementary Fig 2A**). After applying QC filters (**Supplementary Fig 2B**), unsupervised clustering revealed eight major cell types (**Fig 2A**, **Supplementary Fig 2C**). Cell identities were annotated using cell-type specific markers (**Supplementary Fig 2D**). The major immune cell clusters identified were myeloid cells and T/NK-cells. Smaller populations of B-cells and neutrophils were also detected. Further sub-clustering of myeloid cells based on their transcriptomic features demonstrated five myeloid subpopulations (**Fig 2B-C**). In addition to monocytes (Mono) and dendritic cells (DC), three groups of macrophages (defined by the expression of canonical macrophage genes such as *Cd68*, *C1qa*, *C1qb*, *C1qc*, **Supplementary Fig 3**) were identified. Analysis of individual marker genes (**Fig 2D**) and marker gene scores (**Fig 2E**) demonstrated that cardiac resident macrophages (*Cd163*^+^ resident Mac) expressed canonical markers of tissue resident macrophages including *Cd163*, *Lyve1*, *Folr2*, and *Cbr2* ^23, 24^. The other two macrophage subpopulations, *Cxcl9*^+^*Cxcl10*^+^ Mac and *Nlrp3*^+^ Mac each expressed *Ccr2* and were differentiated by specific marker gene signatures (*Cxcl9*^+^*Cxcl10*^+^ Mac: *Cxcl9*, *Cxcl10*) and (*Nlrp3*^+^ Mac: *Nlrp3*, *Ccl4*, *Cd14*). Among these populations, the abundance of *Cxcl9^+^Cxcl10^+^* macrophages increased in *Ctla4^+/-^Pdcd1^-/-^* hearts compared to *Ctla4^+/-^Pdcd1^-/-^*hearts (**Fig 2C**). We further generated single cell RNA-seq data from wide type (WT) mouse hearts (n=6) and mapped the dataset onto the original object consisting of *Ctla4^+/+^Pdcd1^-/-^* (n=4) and *Ctla4^+/-^ Pdcd1^-/-^* (n=10) myeloid cells. Myeloid cells in WT mice predominantly mapped to 3 subclusters: *Cd163* resident Mac, DCs, and monocytes with prediction scores of greater than 0.75 (**Supplementary Fig 4**). In contrast, very few cells mapped to *Cxcl9 Cxcl10* macrophages. Cells that did map to the *Cxcl9 Cxcl10* Mac subset displayed very low prediction scores. These data indicate that *Cxcl9 Cxcl10* macrophages cannot be confidently identified within the query (WT) dataset (**Supplementary Fig 4C, D**).

**Fig 2.**
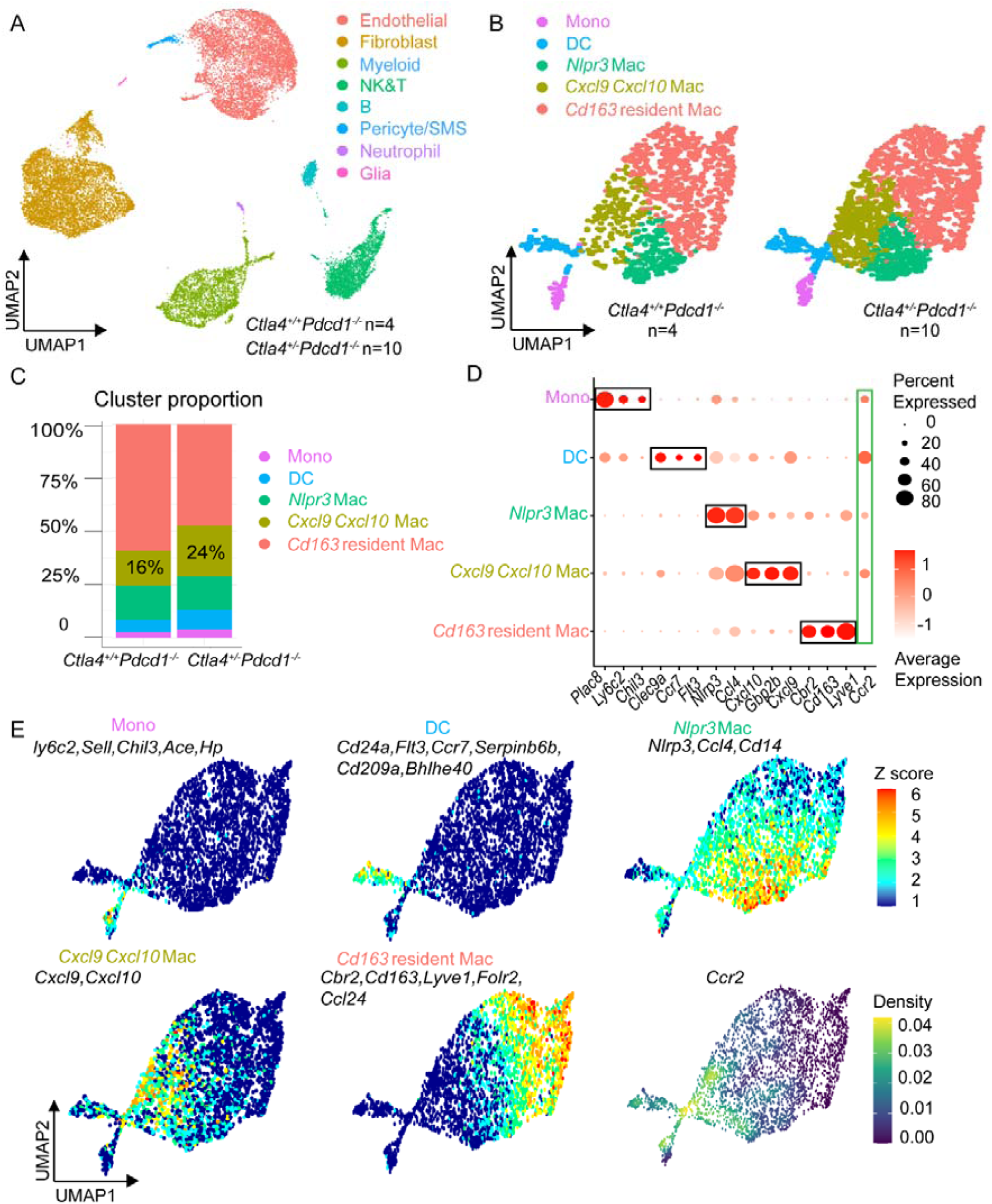
Expansion of *Cxcl9^+^Cxcl10^+^* macrophages in *Ctla4^+/-^Pdcd1^-/-^*mouse hearts. (A) UMAP clustering of 23,606 cells from 14 mouse hearts (*Ctla4^+/+^Pdcd1^-/-^*, n=4; *Ctla4^+/-^Pdcd1^-/-^*, n=10), showing 8 major cell types. (B) UMAP clustering of 3,209 the myeloid cells spilt by experimental group highlighting 5 transcriptionally distinct subclusters. (C) The proportion of each myeloid subcluster in *Ctla4^+/+^Pdcd1^-/-^* and *Ctla4^+/-^Pdcd1^-/-^*mice. (D) Dot plots of differentially expressed genes in each myeloid subcluster. (E) Z-score feature plot of enriched genes in each myeloid subcluster and density plot of *Ccr2* expression. Cell state marker genes (in black) were selected based on robust enrichment in their respective subclusters.

### *Cxcl9^+^Cxcl10^+^* macrophages exhibited an activated phenotype in ICI myocarditis mouse hearts

To further investigate differences between myeloid cells found within *Ctla4^+/+^Pdcd1^-/-^* and *Ctla4^+/-^Pdcd1^-/-^* hearts, we performed a differential gene expression analysis using our scRNA-seq dataset. *Cxcl9*, *Cxcl10*, *Gbp2b*, *Ccl8*, Ccl5, *Fcgr4*, *Ly6a*, *Lgals3*, and *AW112010* were upregulated in myeloid cells from *Ctla4^+/-^Pdcd1^-/-^* mice (**Fig 3A**, **Supplementary Fig 5).** The top 10 upregulated genes in *Ctla4^+/-^Pdcd1^-/-^*mice were exclusively expressed in the *Cxcl9^+^Cxcl10^+^*macrophage subcluster (**Fig 3B**) suggesting that the expansion and activation of *Cxcl9*^+^*Cxcl10*^+^ macrophages represented the predominate difference in myeloid cells between experimental groups. Further validation of expression of *Cxcl9, Cxcl10, Gbp2b*, *Ccl8,* and *Fcgr4* mRNA expression using RT-PCR revealed striking increases in the myocardium of *Ctla4^+/-^Pdcd1^-/-^*compared to *Ctla4^+/+^Pdcd1^-/-^* hearts (**Fig 3C**). *In situ* hybridization revealed increased absolute abundance frequency of *Cxcl9^+^Ccr2^+^*and *Cxcl10^+^Ccr2^+^* macrophages in *Ctla4^+/-^Pdcd1^-/-^* hearts compared with wide type and *Ctla4^+/+^Pdcd1^-/-^*hearts. The majority of *Cxcl9^+^* and *Cxcl10^+^* cells co-expressed CCR2 and were identified as macrophages in *Ctla4^+/-^Pdcd1^-/-^* hearts (**Fig 3D**). Flow cytometry showed increased FCGR4 protein expression on *Ctla4^+/-^Pdcd1^-/-^* macrophages (**Fig 3E)**. To identify *Cxcl9^+^Cxcl10*^+^ macrophages subsets among CCR2^+^ macrophages by flow cytometry, we examined the expression of FCGR4 (a surface protein). CCR2^-^ macrophages did not express FCGR4 in Ctla*4^+/+^Pdcd1^-/-^* hearts. We observed significantly increased frequency of CCR2^+^FCGR4^high^ macrophages in *Ctla4^+/+^Pdcd1^-/-^* hearts compared with *Ctla4^+/+^Pdcd1^-/-^* hearts (**Supplementary Fig 18**).

**Fig 3.**
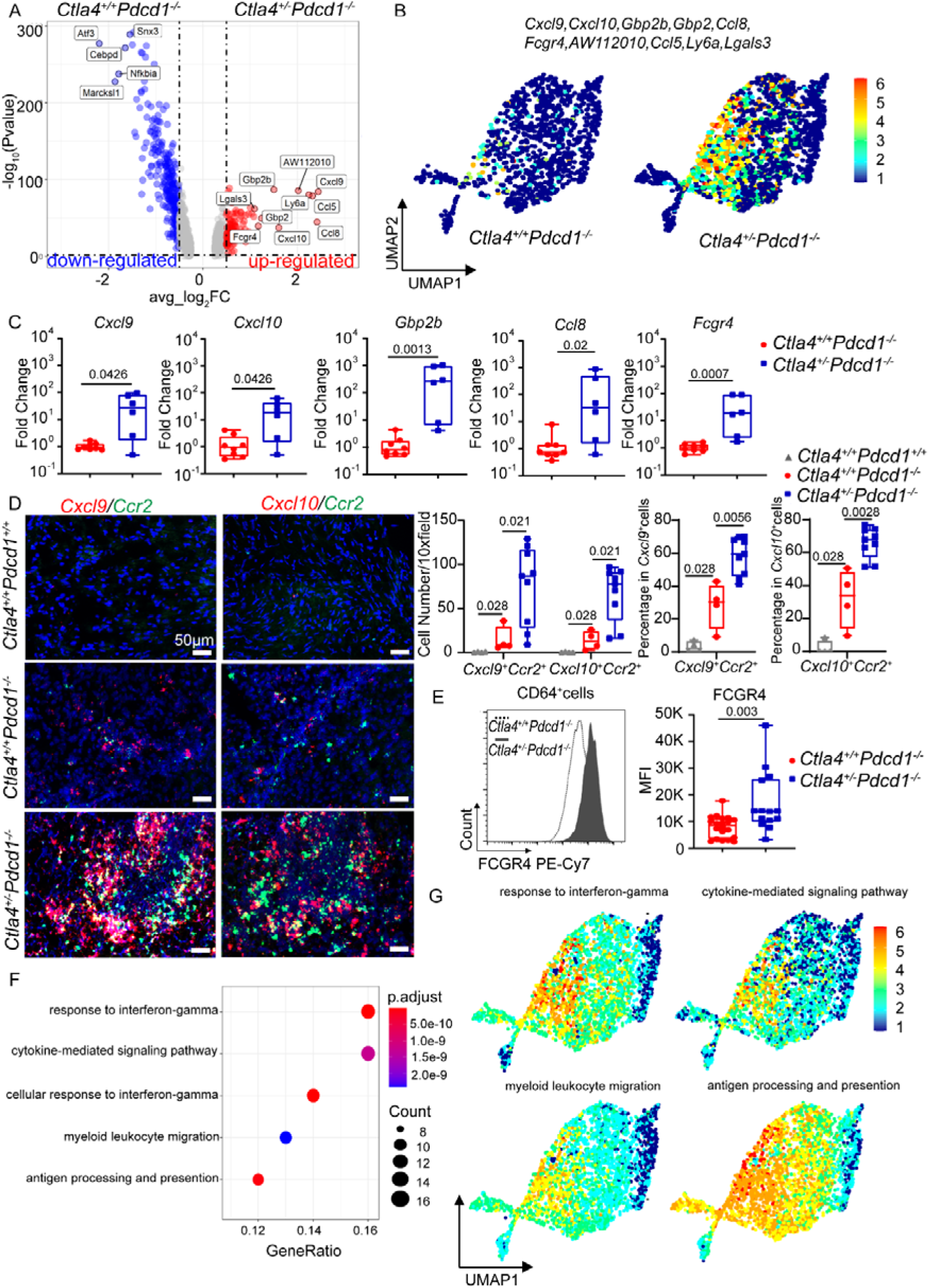
*Cxcl9^+^Cxcl10^+^*macrophages exhibit an activated phenotype in ICI myocarditis. (A) Volcano plot of differentially expressed genes amongst myeloid cells from *Ctla4^+/+^ Pdcd1^-/-^ and Ctla4^+/-^Pdcd1^-/-^*hearts obtained by Wilcoxon Rank Sum test using R package Seurat (v4). (B) Z-score feature plot of the top 10 up-regulated genes in *Ctla4^+/+^ Pdcd1^-/-^* myeloid cells compared to *Ctla4^+/-^Pdcd1^-/-^*myeloid cells split by experimental group. Differentially expressed genes are selectively expressed in *Cxcl9^+^Cxcl10^+^* macrophages (C) Increased *Cxcl9, Cxcl10, Gbp2b, Ccl8,* and *Fcgr4* mRNA expression in *Ctla4^+/+^ Pdcd1^-/-^* compared to *Ctla4^+/-^ Pdcd1^-/-^* heart tissue measured by RT-PCR. Data collected from two independent experiments, *Ctla4^+/+^Pdcd1^-/-^* (n=8), *Ctla4^+/-^Pdcd1^-/-^* (n=6), Mann-Whitney test, two-tailed. (D) Co-expression of *Cxcl9 and Cxcl10* with *Ccr2* in mouse hearts visualized by RNA in situ hybridization. (Left) Representative images in each condition, scale bar, 50 µm. (Right) Quantification of the cell number of *Cxcl9^+^Ccr2^+^*cells or *Cxcl10^+^Ccr2^+^*cells per 10X field in each condition as well as the percentage of *Cxcl9^+^Ccr2^+^* or *Cxcl10^+^Ccr2^+^*cells in *Cxcl9^+^* or *Cxcl10^+^*cells. *Ctla4^+/+^Pdcd1^+/+^*(n=4), *Ctla4^+/+^Pdcd1^-/-^* (n=4), *Ctla4^+/-^Pdcd1^-/-^* (n=9), Mann-Whitney test, two-tailed. (E) Quantification of FCGR4 protein expression on CD64^+^ macrophages by flow cytometry. Data collected from four independent experiments, *Ctla4^+/+^Pdcd1^-/-^* (n=20), *Ctla4^+/-^Pdcd1^-/-^* (n=14), Mann-Whitney test, two-tailed. (F) GO pathway enrichment analysis of up-regulated genes in *Ctla4^+/-^Pdcd1^-/-^* myeloid cells. The top five enriched pathways in *Ctla4^+/-^Pdcd1^-/-^* myeloid cells are displayed. Genes used in the analysis were selected from Seurat differential expression with P < 0.05 and log_2_FC > 0.5. P value calculated using hypergeometric distribution and corrected for multiple comparisons. (G) Z-score feature plots of enriched genes involving in response to Interferon-gamma; cytokine-mediated signaling pathway; myeloid leukocyte migration; antigen processing and presentation pathways in myeloid cells.

GO (Gene Ontology) enrichment pathway analysis using genes up-regulated in *Ctla4^+/-^Pdcd1^-/-^* mice indicated that response to IFN-γ signaling, cytokine mediated signaling, myeloid cell migration, and antigen presentation were among the most impacted pathways (**Fig 3F**). Each of these pathways localized to the *Cxcl9*^+^*Cxcl10*^+^ macrophage cluster (**Fig 3G**). These data suggest that *Cxcl9^+^Cxcl10^+^* macrophages in ICI myocarditis may represent an activated population with enhanced potential for migration, inflammatory cytokine/chemokine production, IFN-γ signaling, and antigen presentation. We also observed a similar phenotype in male *Ctla4^+/-^Pdcd1^-/-^*hearts demonstrated by increased CD68^+^ macrophage accumulation and expansion of *Cxcl9^+^Cxcl10^+^* macrophages compared with male *Ctla4^+/+^Pdcd1^-/-^* hearts (**Supplementary Fig 6**). In addition, *Cxcl9^+^Cxcl10^+^* macrophages exhibited distinct phenotypes compared with *Nlrp3^+^* macrophages or *Cd163^+^* resident macrophages (**Supplementary Fig 7**). Compared with *Cxcl9^+^Cxcl10^+^* macrophages, *Nlrp3*^+^ macrophages exhibited enrichment for genes implicated in the response to LPS, regulation of IL-1β production, and stromal cell proliferation. While *Cd163^+^*resident macrophages were displayed enrichment for pathways involved in regulating epithelial cell proliferation and response to stress.

Furthermore, gene regulatory network analysis using the SCENIC (single-cell regulatory network inference and clustering) pipeline, which identifies modules of transcription factors co-expressed with their target genes, referred to as regulons^25^, revealed that activated *Cxcl9*^+^*Cxcl10*^+^ macrophage in ICI myocarditis showed a unique and specific transcriptional regulatory network. Downstream transcription factors of IFN-γ including STAT1 and IRF7 were enriched in activated *Cxcl9*^+^*Cxcl10*^+^ macrophage (**Supplementary Fig 8A**). RNA transcripts of Stat1 and Irf7 were present in the *Cxcl9*^+^*Cxcl10*^+^ macrophage subcluster (**Supplementary Fig 8B**). TF regulatory network analysis predicted several downstream genes regulated by these TFs in the *Cxcl9*^+^*Cxcl10*^+^ macrophage subcluster, including *Cxcl9, Cxcl10, Gbp2, Fcgr4* (**Supplementary Fig 8C**). To validate this, we analyzed the expression and phosphorylated state of STAT1 protein in hearts from *Ctla4^+/+^Pdcd1^-/-^* and *Ctla4^+/-^ Pdcd1^-/-^* mice (**Supplementary Fig 8D**). Compared with control *Ctla4^+/+^Pdcd1^-/-^* mice, *Ctla4^+/-^Pdcd1^-/-^* mice exhibited significant enhanced expression and phosphorylation levels of STAT1, indicating the activation of IFN-γ signaling pathway. *In vitro*, we also found that inhibition of STAT1 activation using a JAK1/2 inhibitor Ruxolitinib significantly suppressed IFN-γ induced *Cxcl9* expression in BMDMs (**Supplementary Fig 8E**). Knockdown of *Stat1* expression in BMDMs by siRNA significantly reduced IFN-γ induced Cxcl9 expression (**Supplementary Fig 8F**). These findings indicate an important role for IFN-γ-STAT1 signaling in the emergence of *Cxcl9^+^Cxcl10^+^*macrophages.

### *Cxcl9^+^Cxcl10^+^* macrophages originated from CCR2^+^ monocytes

To gain further insights into the relationships between myeloid subcluster, we performed trajectory analysis using Palantir^26^. We specified the starting point as monocytes. Calculation of pseudotime values suggested that *Cd163*^+^ macrophages, *Cxcl9^+^Cxcl10^+^* macrophages, *Nlrp3^+^* macrophages, and DCs represented differentiated cell states. Ascertainment of differentiation potential (entropy) values indicated that *Cxcl9^+^Cxcl10^+^* macrophages displayed the lowest level of cell plasticity highlighting that this may represent a highly specialized cell state (**Fig 4A-B, D**). We then calculated terminal state probabilities for DCs, *Cd163*^+^ macrophages, and *Cxcl9^+^Cxcl10^+^* macrophages differentiating from monocytes. DCs and *Cd163*^+^ macrophage terminal state probabilities were largely restricted to cells within their respective clusters. In contrast, high terminal state probabilities for the *Cxcl9^+^Cxcl10^+^*macrophage state was evident suggesting a development relationship from monocytes (**Fig 4C-D**). Compared with control mice, *Cxcl9^+^Cxcl10^+^*macrophages from *Ctla4^+/+^Pdcd1^-/-^* mice exhibited lower entropy and higher terminal state probability values suggesting these macrophages in disease mice are more differentiated and functionally activated (**Fig 4D**). In addition, we employed a neutralizing CCR2 antibody (MC-21) to inhibit monocyte mobilization in *Ctla4^+/-^ Pdcd1^-/-^* mice. MC-21 blocks the interaction between CCR2 and CCL2 which prevents the recruitment of CCR2^+^ monocytes to sites of inflammation and results in reduction of macrophages derived from recruited CCR2^+^ monocytes ^27, 28^. Consistent with the findings of our trajectory analysis, MC-21 antibody treatment reduced the accumulation of CD68^+^ macrophages (**Supplementary Fig 9**), the abundance of Cxcl9^+^Cxcl10^+^ macrophages (**Fig 4E-F**), and reduced frequency of CCR2^+^FCGR4^high^ macrophages (**Supplementary Fig 18**). MC-21 blocks the interaction between CCR2 and CCL2 and prevents the recruitment of CCR2^+^ monocytes to sites of inflammation^27, 28^. Through lineage tracing of recruited monocytes, we have confirmed that MC-21 displays similar properties in the heart (**Supplementary Fig 10**). These findings demonstrate the dependence of *Cxcl9^+^Cxcl10^+^*macrophages on monocytes.

**Fig 4.**
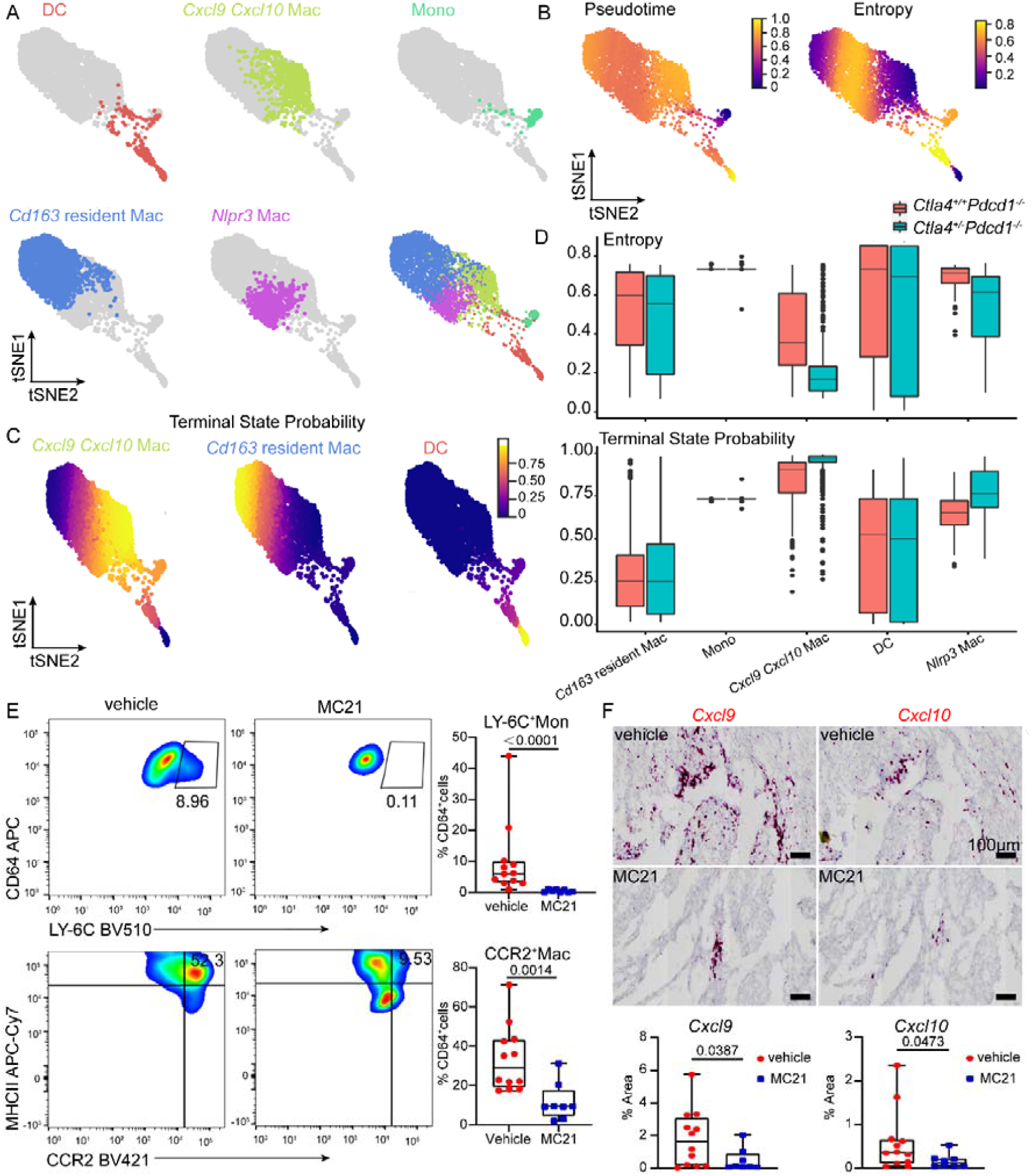
*Cxcl9^+^Cxcl10^+^*macrophages originate from monocytes. (A) tSNE force-directed layout plot of myeloid cells. Cells are colored by cell cluster annotations. (B) Pseudotime and entropy values of myeloid cells. *Cxcl9^+^Cxcl10^+^*macrophages (*Cxcl9 Cxcl10* Mac) have high pseudotime and low entropy values suggesting that they represent a differentiated cell state. (C) Terminal State Probability of cell states predicted as differentiated populations: *Cxcl9 Cxcl10* Mac; *Cd163* resident Mac; and DCs. (D) Box plots of entropy (upper) and *Cxcl9 Cxcl10* Mac terminal state probability (lower) of myeloid subclusters split by experimental group. (E) Percentage of LY-6C^high^ monocytes and CCR2^+^ macrophages of cardiac CD64^+^ cells from vehicle or MC-21 antibody treated mice quantified by flow cytometry. Displayed cells are CD45^+^LY-6G^-^CD64^+^. Data collected from four independent experiments. vehicle group (n=12), MC-21 *treated group* (n=8), Mann-Whitney test, two-tailed. (F) Representative images (upper) and quantification (lower) of *Cxcl9 and Cxcl10* positive cells in the heart 6 days after MC-21 antibody treatment. Data collected from four independent experiments. vehicle group (n=12), MC-21 *treated group* (n=8), Mann-Whitney test, two-tailed.

### *CXCL9^+^CXCL10*^+^ macrophages in clinical cases of ICI myocarditis

We next examined whether macrophages expressing *CXCL9*, *CXCL10*, and CD16α (human homologue of mouse FcgR4) were present with the myocardium of patients with ICI myocarditis. *In situ* hybridization revealed robust expression of *CXCL9* and *CXCL10* mRNA in interstitial cells within the myocardium of patients with biopsy conformed ICI myocarditis. In contrast, rare cells expressing *CXCL9* and *CXCL10* mRNA were detected in healthy donors, and subjects with dilated cardiomyopathy (DCM), ischemic cardiomyopathy (ICM), and lymphocytic myocarditis (LM) (**Fig 5A**). Immunostaining further revealed the presence of CD16α^+^ macrophages selectively expanded in subjects with ICI myocarditis. Consistent with our mouse scRNA-seq data, CD16α^+^ macrophages expressed CCR2 in ICI myocarditis samples. Few CD16α^+^ macrophages observed in DCM, ICM, and LM (**Fig 5B).** *In situ* hybridization and immunostaining performed on consecutive sections obtained from ICI myocarditis specimens revealed co-location of *CXCL9* and *CXCL10* mRNA with CD68, CD16α, and CCR2 protein (**Supplementary Fig 11**). These findings confirm the existence of *CXCL9^+^CXCL10^+^* macrophages in human ICI myocarditis.

**Fig 5.**
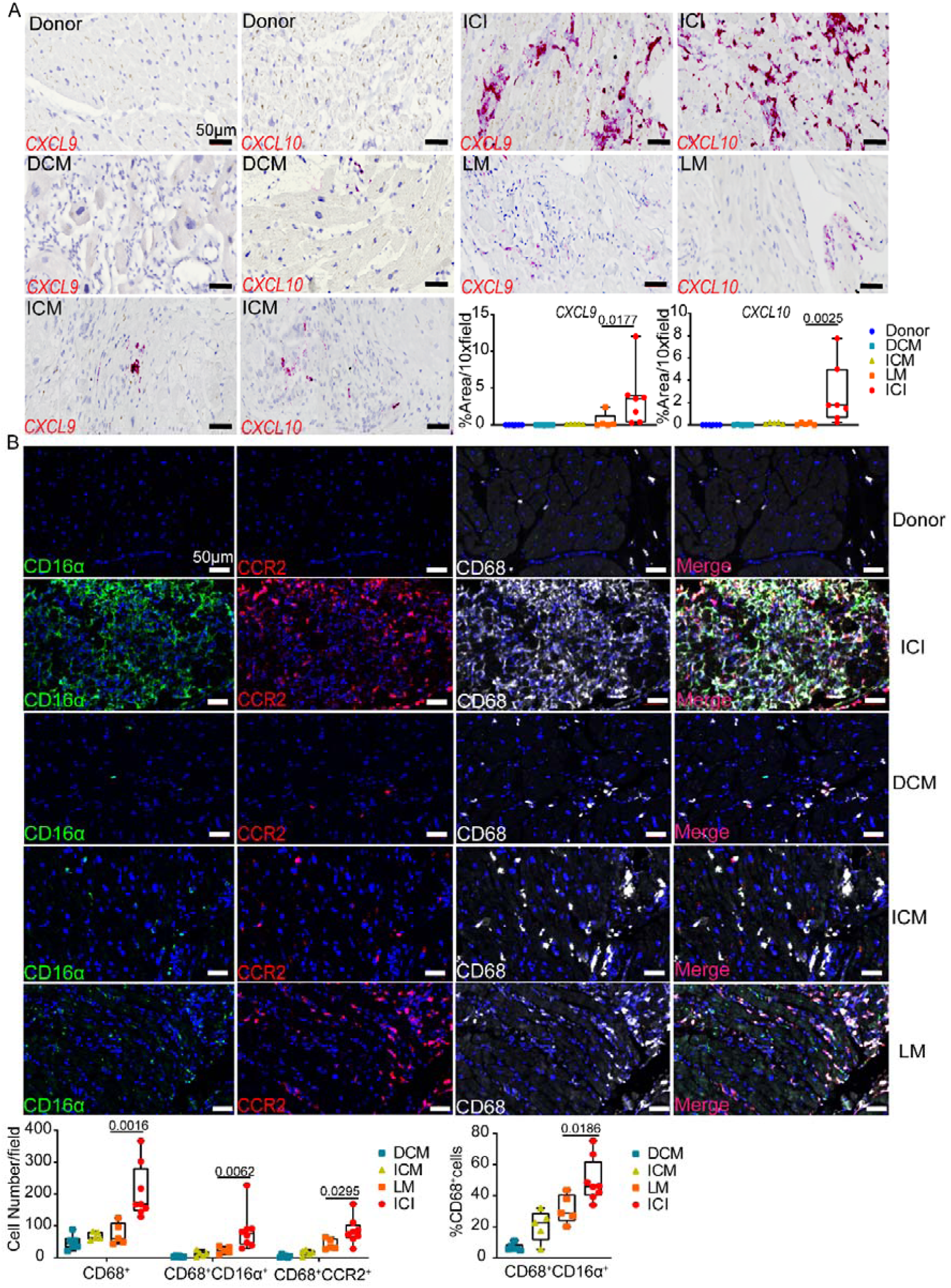
*CXCL9^+^CXCL10^+^ macrophages in* human ICI associated myocarditis. (A) Expression of *CXCL9* and *CXCL10* via RNA in situ hybridization in human heart tissue from patients with ICI myocarditis (ICI, n=7), lymphocytic myocarditis (LM, n=5), ischemic cardiomyopathy (ICM, n=5), dilated cardiomyopathy (DCM, n=6), and donor control subject (n=6). Quantification of the number of *CXCL9^+^* and *CXCL10^+^* cells, Mann-Whitney test, two-tailed. Scale bar 50 µm. (B) Immunofluorescent staining of CD16α (green), CCR2 (red), CD68 (white) and DAPI (blue) in human heart tissue from patients with ICI (n=8), LM (n=5), ICM (n=5), DCM (n=6) and donor control subjects (n=6). Quantification of cell number and the percentage of CD68^+^CD16a^+^ cells in all CD68^+^cells, Mann-Whitney test, two-tailed. Scale bar, 50 µm.

### T-cells are the major source of IFN-γ in ICI myocarditis mouse hearts

Pathway analysis indicated that *Cxcl9^+^Cxcl10^+^*macrophages express an IFN-γ activation signature suggesting that cardiac IFN-γ signaling may regulate the differentiation or activity of this macrophage population. Consistent with this possibility, we detected increased expression of *Ifng* mRNA in *Ctla4^+/-^Pdcd1^-/-^* hearts compared to *Ctla4^+/+^Pdcd1^-/-^* control hearts (**Fig 6A**). To identify the cellular source of IFN-γ, we leveraged our scRNA-seq dataset and observed robust expression of *Ifng* mRNA in the T/NK-cell cluster (**Fig 6B**). High-resolution clustering of T-cells and NK-cells identified four major subclusters including CD8^+^ T-cells, CD4^+^ T-cells, NK-cells and naïve T-cells with differing transcriptional signatures (**Supplementary Fig 12A**). *Ifng* mRNA was readily expressed in CD4^+^ and CD8^+^ T-cells (**Fig 6C**). Intracellular flow cytometry confirmed that CD4^+^ and CD8^+^ T-cells produced increased amounts of IFN-γ in *Ctla4^+/-^Pdcd1^-/-^* hearts compared to controls. Macrophages and NK-cells produced low amounts of IFN-γ. Among T-cell populations, CD8^+^ T-cells displayed the greatest increase in IFN-γ expression (**Fig. 6D**).

**Fig 6.**
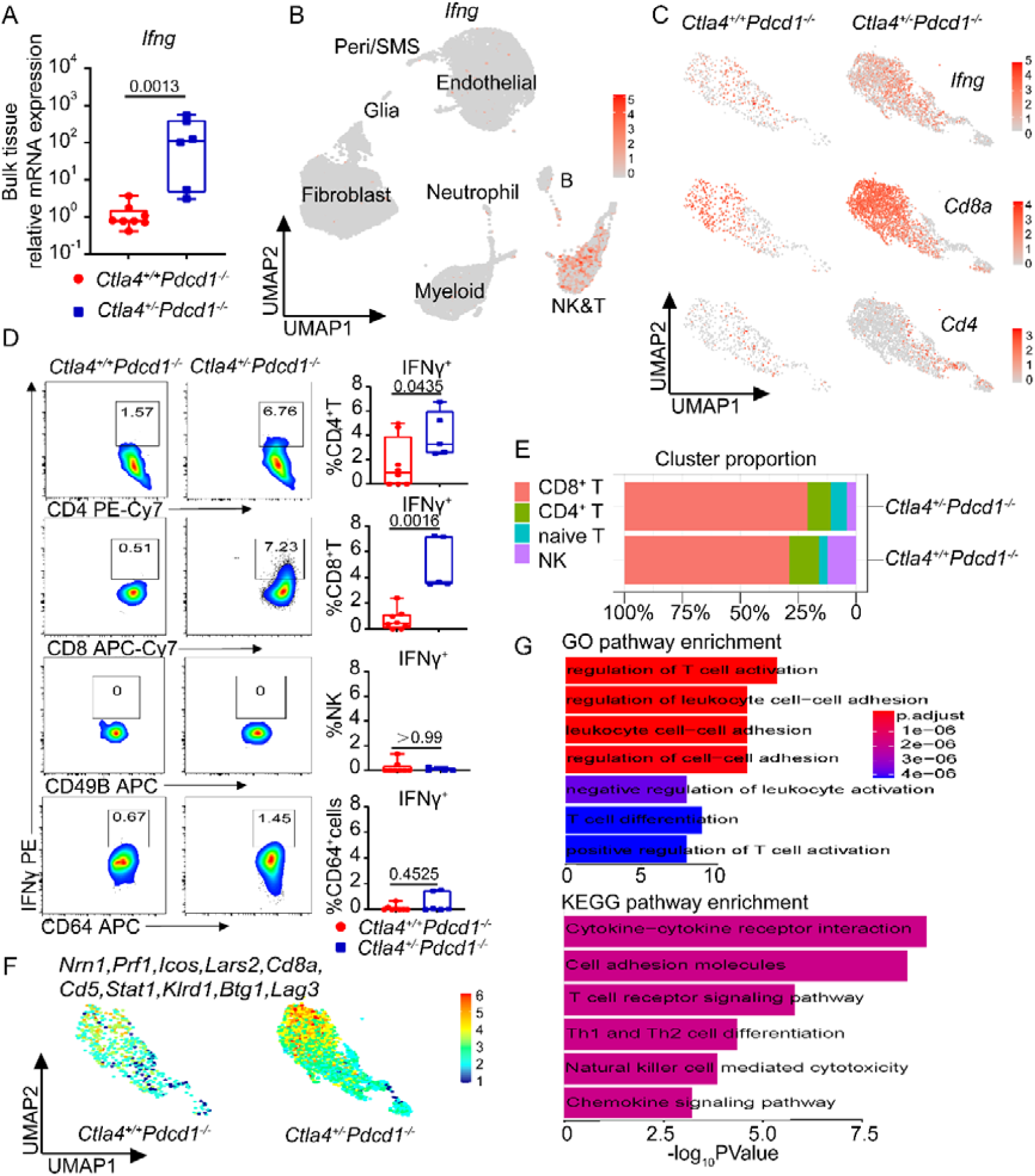
T-cells are the primary source of IFN-γ in ICI myocarditis mouse hearts. (A) Increased *Ifng mRNA* expression in *Ctla-4^+/-^Pdcd1^-/-^* mouse hearts measured by RT-PCR. Data collected from two independent experiments, *Ctla4^+/+^Pdcd1^-/-^* (n=8), *Ctla4^+/-^Pdcd1^-/-^*(n=6), Mann-Whitney test, two-tailed. (B) Feature plot of *Ifng* expression in all cell types recovered from the heart showing specific expression in the NK/T-cell cluster. (C) Feature plots of *Ifng, Cd8a,* and *Cd4* expression in NK&T-cells showing CD8 T-cell expansion and enriched *Ifng* expression in CD8 T-cells from *Ctla4^+/-^Pdcd1^-/-^*hearts. (D) Percentages of IFNγ^+^CD4^+^, IFNγ^+^CD8^+^ T-cells, IFNγ^+^NK-cells, and IFNγ^+^CD64^+^ macrophages analyzed by flow cytometry. Data collected from two independent experiments, *Ctla4^+/+^Pdcd1^-/-^* (n=8), *Ctla4^+/-^Pdcd1^-/-^*(n=5), Mann-Whitney test, two-tailed. (E) The proportion of each NK&T subcluster per experimental group. (F) Z-score feature plots of top 10 up-regulated genes in *Ctla4^+/-^Pdcd1^-/-^* NK&T-cells compared to *Ctla4^+/+^Pdcd1^-/-^* NK&T cells split by group.(G) GO and KEGG enriched pathways using genes up-regulated in *Ctla4^+/-^Pdcd1^-/-^*NK&T cells compared to *Ctla4^+/+^Pdcd1^-/-^* NK&T cells. Genes used in the analysis were selected from Seurat differential expression with P < 0.05 and log_2_FC > 1. P-values calculated by hypergeometric distribution using R package ClusterProfiler.

Detailed examination of T-cell subclusters revealed expansion of CD8^+^ T-cells in *Ctla4^+/-^Pdcd1^-/-^* hearts compared to *Ctla4^+/+^Pdcd1^-/-^* control hearts (**Fig 6E**). Differential expression analysis identified robust transcriptional differences between experimental groups. Top 10 genes up-regulated in *Ctla4^+/-^Pdcd1^-/-^* T/NK-cells (*Nrn*1, *Prf1*, *Icos*, *Lars2*, *Cd8a*, *Cd5*, *Stat1*, *Klrd1*, *Btg1*, *Lag*3) mapped to CD8^+^ T-cells (**Fig 6F**). GO and KEGG pathway analysis implicated enhanced T-cell activation, T-cell differentiation and cytokine-cytokine receptor signaling as putative markers of T-cell activation in *Ctla4^+/-^Pdcd1^-/-^* hearts (**Fig 6G, Supplementary Fig 12B-C**). These data highlight the putative pathological role of CD8^+^ T-cells in our mouse model of ICI myocarditis.

### IFN-γ and CXCR3 signaling are predicted to mediate crosstalk between *Cxcl9^+^Cxcl10^+^* macrophages and T-cells

To investigate crosstalk between macrophages and T-cells in ICI myocarditis, we applied CellChat to infer cell-cell communication. We identified several possible ligand-receptor interactions. Predicted signals outgoing from CD4^+^ T-cells and CD8^+^ T-cells included IFN-II, MIF, and FASL. Each of these signals was predicted to be received by *Cxcl9^+^Cxcl10^+^* macrophages. CXCL ligands and CD137 were predicted to signal from *Cxcl9^+^Cxcl10^+^* macrophages to T-cells (**Supplementary Fig 13**).

Among these pathways, we focused on IFN-II and CXCL signaling given their enrichment in *Cxcl9^+^Cxcl10^+^* macrophages and CD8^+^ T-cells, respectively. We first examined *Ifng*, *Ifngr1* and *Ifngr2* mRNA expression in macrophages and T-cells. *Ifng* was selectively expressed in CD4^+^ and CD8^+^ T-cells. *Ifngr1* was expressed in all macrophages and T-cells and *Ifngr2* was exclusively expressed in *Cxcl9^+^Cxcl10^+^*macrophages (**Fig 7A)**. CellChat pathway network analysis predicted that CD4^+^ T-cell and CD8^+^ T-cells were the primary ligand sources and that *Cxcl9^+^Cxcl10^+^*macrophages served as the receivers of IFN-γ signaling (**Fig 7B-C**). CD8^+^T-cells demonstrated a higher communication probability compared to CD4^+^ T-cells (**Fig 7D**).

**Fig 7.**
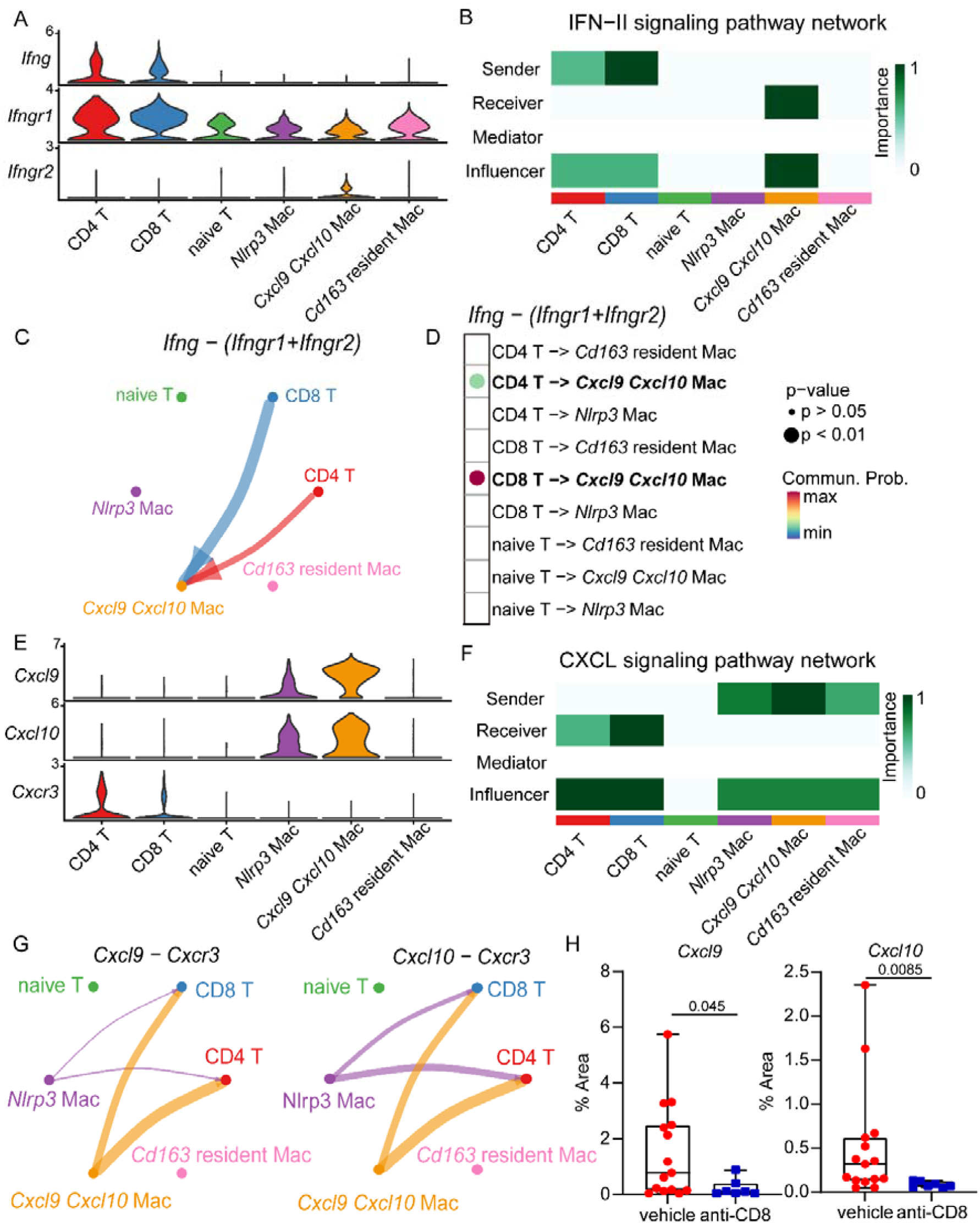
T-cells are predicted to orchestrate the expansion and activation of *Cxcl9^+^Cxcl10^+^* macrophages. (A) Cell to cell communication analysis using CellChat predicted that T-cells signal to macrophages through IFN-γ. Violin plot showing the expression distribution of IFN-γ pathway ligand and receptors in T-cells and macrophages. (B) Heatmap showing the relative importance of each cell state based on the computed network of IFN-γ signaling. (C) Circle plot summarizing the inferred intercellular communication network between T-cells and macrophages for IFN-γ signaling. (D) Dotplot showing the strength of interaction between T-cell and macrophage cell states for IFN-γ signaling. P values were calculated using R package CellChat. (E) Violin plot showing the expression distribution of signaling genes involved in the inferred reciprocal CXCL signaling network (Cxcl9/ Cxcl10-Cxcr3) between macrophages and T-cells. (F) Heatmap displaying the relative importance of each cell state based on the computed network of CXCL signaling. (G) Circle plot depicting the inferred intercellular communication network between macrophage and T-cell states for CXCL signaling (Cxcl9/ Cxcl10-Cxcr3). (H) Quantification of cardiac *Cxcl9^+^*and *Cxcl10^+^ cells* 6 days after anti-CD8 antibody treatment. Data collected from three independent experiments. vehicle (n=15); anti-CD8 (n=7), Mann-Whitney test, two-tailed.

Reciprocal signaling from *Cxcl9^+^Cxcl10^+^* macrophages to T-cells was further explored using CellChat, which predicted that these cell types interact through CXCL signaling. *Cxcl9^+^Cxcl10^+^* macrophages were predicted to serve as the source of CXCL ligands that signaled to CD4^+^T-cell and CD8^+^T-cells through the CXCR3 receptor (**Fig 7E-G**). CXCR3 is reported to activate mitogen-activated protein kinases and phosphoinositide 3-kinase/protein kinase B (AKT) pathways leading to activation, differentiation, and recruitment of T-cells^29, 30^. Cellchat also identified CXCL16-CXCR6 signaling as a second pathway that may mediate interactions between macrophages and T-cells (**Supplemental Fig 14**).

To evaluate the predicted interaction between CD8^+^ T-cells and *Cxcl9^+^Cxcl10^+^* macrophages, we depleted CD8^+^ T-cells in *Ctla4^+/-^Pdcd1^-/-^* mice with anti-CD8 antibody beginning at 4 weeks of life (**Supplementary Fig 15A**). CD8 depletion reduced *Cxcl9^+^Cxcl10^+^* macrophages in the heart indicated by decreased cardiac *Cxcl9* and *Cxcl10* expression (**Supplementary Fig 15B,** **Fig 7H**) and reduced frequency of CCR2^+^FCGR4^high^ macrophages (**Supplementary Fig 18**). We also leveraged the bulk RNA-seq dataset from 9 human ICI associated myocarditis biopsy samples^21^ and observed a positive correlation between *CD8A* and *CXCL9/10/FCGR3A* expression (**Supplementary Fig 16**). Collectively, these findings support a role for CD8^+^ T-cells in the expansion and activation of *Cxcl9^+^Cxcl10^+^*macrophages.

### IFN-γ blockade and macrophage depletion reduced *Cxcl9^+^Cxcl10^+^* **macrophages and prolonged survival of *Ctla4^+/-^Pdcd1^-/-^mice*.**

To examine the causal role of IFN-γ signaling in the activation and expansion of *Cxcl9^+^Cxcl10^+^* macrophages and impact on ICI myocarditis, we examined the effects of blocking IFN-γ signaling. *Ctla4^+/-^Pdcd1^-/-^*mice were treated with either isotype control or IFN-γ neutralization antibody (Clone, R46A2) beginning at 3 weeks of life. IFN-γ blockade significantly prolonged the survival of *Ctla4^+/-^Pdcd1^-/-^* mice (**Fig 8A**). Previous studies have demonstrated that these mice die from myocarditis and arrhythmic events^8^. *In situ* hybridization revealed markedly diminished cardiac *Cxcl9* and *Cxcl10* expression in anti-IFN-γ antibody treated *Ctla4^+/-^Pdcd1^-/-^* mice compared with isotype control treated *Ctla4^+/-^Pdcd1^-/-^* mice, implying a reduction in *Cxcl9^+^Cxcl10^+^* macrophages (**Fig 8B**). IFN-γ blockade reduced the frequency of CCR2^+^FCGR4^high^ macrophages in the heart consistent with a reduction in *Cxcl9^+^Cxcl10^+^*macrophages (**Supplementary Fig 18**). Within the CD64^+^ compartment, we observed that CCR2^+^ cells consisted of CCR2^+^MHC^high^ macrophages and CCR2^+^MHC^low^ monocytes/macrophages (CCR2^+^ monocytes and differentiating macrophages^16^). We observed reduced percentage and absolute number of CCR2^+^MHC^low^ monocytes/macrophages in the heart after 3 weeks of IFN-γ blockade. We did not detect any differences in the abundance of CCR2^+^MHC^high^ macrophages in the heart. These data indicate that IFN-γ reshapes CCR2^+^MHCII^high^ macrophages towards an activated *Cxcl9^+^Cxcl10^+^* phenotype (*Cxcl9^+^Cxcl10^+^Ccr2^+^*macrophages) and influences the recruitment of CCR2^+^ monocytes and their differentiation into macrophages (**Supplementary Fig 17A**). Importantly, we did not detect significant changes in T-cell abundance and phenotype across CD4 and CD8 subsets between control and anti-IFN-γ treated groups arguing against a direct effect on T-cells at this time point (**Supplementary Fig 17 B-D**). These findings indicate that inhibition of IFN-γ signaling prevented the expansion of activated *Cxcl9^+^Cxcl10^+^* macrophages and delay or prevent the progression of ICI myocarditis. Importantly, temporary blockade of IFN-γ signaling does not interfere with the therapeutic effects of conventional chemotherapy^31, 32^.

**Fig 8.**
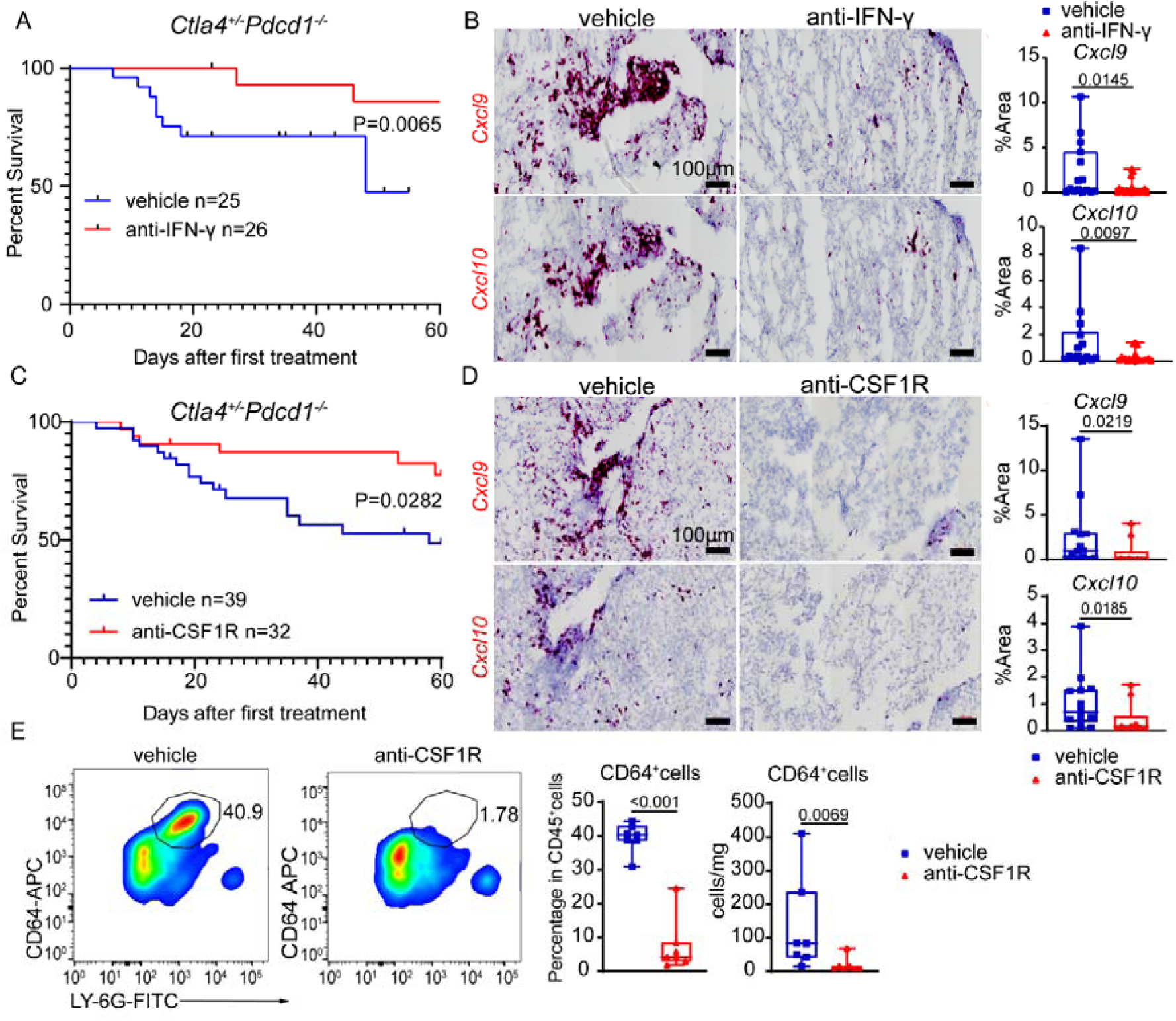
IFN-γ blockade and macrophage depletion *reduce* cardiac *Cxcl9^+^Cxcl10^+^*macrophages and prolong the survival of *Ctla4^+/-^Pdcd1^-/-^ mice*. (A) Survival of *Ctla4^+/-^ Pdcd1^-/-^* mice treated with vehicle (isotype control) or anti-IFN-γ antibody (R46A2). Data collected from four independent experiments vehicle (n=25); anti-IFN-γ (n=26), Log-rank test. (B) Representative images (left) and quantification (right) of cardiac *Cxcl9^+^* and *Cxcl10^+^*cells as determined by RNA in situ hybridization 23 days after vehicle or anti-IFN-γ antibody treatment. Data collected from five independent experiments. vehicle (n=15); anti-IFN-γ (n=21), Mann-Whitney test, two-tailed. (C) Survival of *Ctla4^+/-^ Pdcd1^-/-^* mice treated with vehicle (isotype control) or anti-CSF1R antibody (AFS98). vehicle (n=39); anti-CSF1R (n=32), Log-rank test. (D) Representative images (left) and quantification (right) of cardiac *Cxcl9^+^* and *Cxcl10^+^* cells as determined by RNA in situ hybridization 23 days after vehicle or anti-CSF1R antibody treatment. Data collected from four independent experiments. vehicle (n=14); anti-CSF1R (n=10), Mann-Whitney test, two-tailed. (E) Cardiac CD64^+^macrophages depletion was verified by flow cytometry. Representative images (left) and quantification (right) of CD64*^+^* cells as determined by flow cytometry 60 days after first vehicle or anti-CSF1R antibody treatment. Displayed cells (left) are gated CD45^+^ cells. Data collected from three independent experiments. vehicle (n=7); anti-CSF1R (n=7), Mann-Whitney test, two-tailed.

To further study the role of *Cxcl9^+^Cxcl10^+^*macrophages in ICI myocarditis, we depleted macrophages in the heart by administrating anti-CSF1R antibody in Ctla*4^+/-^ Pdcd1^-/-^* mice beginning at 3 weeks of age (**Fig 8E**). Anti-CSF1R treatment significantly prolonged the survival of *Ctla4^+/-^Pdcd1^-/-^*mice (**Fig 8C**) and markedly reduced the number of *Cxcl9^+^Cxcl10^+^* macrophages. These findings are consistent with what we observed in the IFN-γ blockade experiments (**Fig 8D**) corroborating the functional importance of macrophages in ICI myocarditis.

## Discussion

ICI myocarditis is a serious manifestation of ICI-induced toxicity. Histological analysis of myocardial specimens from ICI myocarditis patients revealed immune infiltrates comprised of CD4^+^ T-cells, CD8^+^ T-cells and macrophages ^4, 8, 9, 33^. Substantial effort has focused on elucidating mechanisms of T-cell infiltration and expansion within the heart ^9–12^ including the recognition of clonal populations of T-cells that recognize antigens co-expressed in the tumor and the heart ^6^. Markedly, less is known regarding macrophage populations in ICI myocarditis. Within this context, the transcriptomic features of macrophages, their inherent heterogeneity, and impact on disease progression remain incompletely explored.

Here, we employed an established genetic ICI myocarditis model (*Ctla4^+/-^Pdcd1^-/-^*mice), which recapitulates clinical features seen in subjects with fulminant ICI myocarditis^8^. Using a compilation of technologies (scRNA-seq, immunostaining, *in situ* hybridization, flow cytometry, antibody neutralization and molecular imaging), we demonstrated that ICI myocarditis is associated with the expansion of a population of CCR2^+^ macrophages (*Cxcl9^+^Cxcl10^+^*macrophages) that expressed a transcriptional signature indicative of enhanced IFN-γ activation, cytokine generation, myeloid cell migration, and antigen presentation. A similar population of macrophages was identified in human cases of ICI myocarditis which expanded uniquely in this pathological entity. *Cxcl9^+^Cxcl10^+^*macrophages originated from CCR2^+^ monocytes. Mechanistically, *Cxcl9^+^Cxcl10^+^* macrophages were predicted to be specified by activated T-cells through IFN-γ signaling and participate in a positive feedback loop with T-cells through CXCR3 signaling. Depleting CD8^+^T-cells, CD64^+^cells and blocking IFN-γ signaling led to reductions in *Cxcl9^+^Cxcl10^+^* macrophages and improved survival in our mouse model of ICI myocarditis. These data provide support for the possibility that IFN-γ signaling may be considered as a potential target for the treatment of ICI associated myocarditis.

CXCL9 and CXCL10 are robustly expressed by macrophages that emerge in the setting of ICI myocarditis (*Ctla4^+/-^Pdcd1^-/-^*mouse model and human ICI myocarditis specimens). CXCL9 expressing macrophages are similarly found in ICI treated patients and those who develop of ICI colitis ^34, 35^. Trajectory analysis predicted that *Cxcl9^+^Cxcl10^+^* macrophages were derived from recruited monocytes, which was validated by selective depletion of Ly6C^hi^ monocytes. This finding is consistent with the repeated observation that inflammatory macrophage populations are derived from monocytes that infiltrate the heart after injury^16, 36, 37^. *In silico* analysis further indicated that IFN-γ stimulation orchestrated the differentiation of infiltrating monocytes or monocyte-derived macrophages into *Cxcl9^+^Cxcl10^+^* macrophages. The functional importance of IFN-γ signaling and *Cxcl9^+^Cxcl10^+^* macrophages was validated through our neutralizing antibody studies. Notably, anecdotal clinical studies have recently reported that CTLA4-fusion protein abatacept combined with the Janus kinase inhibitor Ruxolitinib, which targets IFN-γ/Jak2/Stat1 signaling decreased cardiovascular death among ICI-myocarditis patients^38^.

*Cxcl9^+^Cxcl10^+^* macrophages may contribute to ICI myocarditis pathogenesis through several distinct mechanisms including enhancing T-cell responses, innate immune cell recruitment, and directly aggravating local tissue damage. *Cxcl9^+^Cxcl10^+^*macrophages express several inflammatory mediators involved in T-cell migration and T-cell receptor signaling, highlighting the possibility that they are primed to augment T-cell activation. Consistent with our findings, ICI myocarditis patients display enhanced expression of CXCR3 on T-cells, particularly memory CD8^+^ T-cells^34^. CD8^+^ T-cells serve as the predominant cardiac T-cell subset in ICI myocarditis, display an activated phenotype, and produce IFN-γ. Using the same mouse model of ICI myocarditis, it was observed that CD8^+^ T-cells are clonally expanded, target alpha-myosin as MHC-I restricted autoantigen and are required for disease pathogenesis^12^. Informatic prediction of cell-cell communication and suggested that CD8^+^ T-cells signal to *Cxcl9^+^Cxcl10^+^* macrophages by secreting IFN-γ. This analysis also predicted a reciprocal feedback loop where *Cxcl9^+^Cxcl10^+^* macrophages communicated with CD8^+^ T-cells through CXCL16/CXCR6 signaling and to CD4^+^ and CD8^+^ T-cells through CXCL9/CXCL10-CXCR3 signaling. Thus, *Cxcl9^+^Cxcl10^+^*macrophages may stimulation the recruitment and activation of CD8^+^ T cells and potentially trigger cytotoxic responses against cardiomyocytes. Consistent with this possibility, CXCL9/10-CXCR3 signaling has previously been reported to accelerate pressure overload–induced cardiac dysfunction and stimulate T-cell recruitment to the mouse heart^30^.

In addition to regulating T-cell chemotaxis, *Cxcl9^+^Cxcl10^+^*macrophages may also participate in promoting peripheral monocyte and neutrophil recruitment. *Cxcl9^+^Cxcl10^+^* macrophages represent a subset of CCR2^+^ macrophages and express chemokines that regulate peripheral leukocyte migration and chemotaxis as predicted by pathway analysis. Following cardiac injury, CCR2^+^ monocytes are recruited to the heart by CCR2^+^ macrophages through an MYD88-dependent mechanism via the production of chemokines including CCL2/MCP1 and CCL7/MCP3^16^. This raises the possibility that *Cxcl9^+^Cxcl10^+^* macrophages further enhance myocardial inflammation by recruiting additional peripheral immune cells.

*Cxcl9^+^Cxcl10^+^* macrophages may also contribute to cardiomyocyte injury and cell death by activating effector T-cells or inducing antibody-dependent cytotoxicity (ADCC) and phagocytosis^39^. De novo autoantibodies (including to cardiac antigens) have been reported to develop in ∼20% of ICI-treated melanoma patients^40, 41^. In the presence of anti-cardiac autoantibodies, *Cxcl9^+^Cxcl10^+^*macrophages are ideally positioned to participate in antibody-dependent cytotoxicity (ADCC) and target cell phagocytosis^39^. IFN signaling enhances ADCC function of macrophages ^42^ and FcgR4 (human CD16α) represents an essential surface receptor to initiate the ADCC^39^.

In considering strategies to treat ICI myocarditis, it is important to recognize that selected therapeutic targets should ideally not interfere with ongoing or established tumor control. At present, conflicting data exists regarding whether IFN-γ signaling is essential for maintaining the therapeutic response to ICIs. Administration of ICIs leads to enhanced IFN-γ, CXCL9, and CXCL10 production in the tumor microenvironment, which facilitates entry of lymphocytes that eliminate tumor cells^43^. Intriguingly, IFN-γ production also promotes tumor immunoevasion^44^ and participates in doxorubicin associated cardiotoxicity ^31, 32^. The exact phase of requirement for IFN-γ signaling in maintaining tumor control following ICI therapy is unclear. Moving forward, it will be essential to determine how IFN-γ blockade compares to other proposed ICI myocarditis treatments including CTLA4-Ig (abatacept) ^8, 45^, anti-CD52 (alemtuzumab) ^46^, and anti-TNFα (infliximab) ^47^ in regards to effects on both ICI myocarditis and tumor control.

Our study is not without limitations. We acknowledge that there is no perfect animal model of ICI myocarditis and that *Ctla4^+/-^Pdcd1^-/-^*mice may only recapitulate some aspects of the disease. We further recognize that IFN-γ blockade represents a starting point and that considerable work remains to elucidate the functions of *Cxcl9^+^Cxcl10^+^*macrophages, dissect downstream signaling mechanisms, and defining optimal therapeutic targets. Short-term treatment (3 weeks) of anti-IFN-γ did not significantly affect the abundance of CD8^+^ T-cells. It remains possible that longer-term inhibition of IFN-γ signaling may have an impact on the number of CD8^+^ T-cells. It is also plausible that anti-IFN-γ treatment impacts CD8^+^ T-cell phenotypes and gene expression. This is an important possibility that will be investigated in future studies. Nonetheless, our findings advance the field by identifying a specific and targetable population of macrophages in ICI myocarditis. In conclusion, we demonstrate that IFN-γ signaling triggers the expansion of an inflammatory population of *Cxcl9^+^Cxcl10^+^*macrophages in ICI myocarditis that are positioned to augment T-cell recruitment, T-cell activation, chemokine/cytokine production, and ADCC and that blockade of IFN-γ signaling may be considered as a potential approach that requires further evaluation for the treatment of this devastating condition.

## Author Contributions

P.M., J.M., K.L. conceived and designed the study and composed the manuscript. P.M., K.L. analyzed all data generated in this study. J.M., K.L. supervised the study. P.M., G.F., A.V., managed the mouse colony and identified appropriate animals for each experiment. P.M., J.L., J.Q., L.L., J.A. performed single cell RNA and bulk RNA-seq data analyses and figure generation. P.M., J.Q. performed antibody mediated depletion and neutralization studies. P.M., A.B., G.B. prepared samples for single cell sequencing. G.S.H., H.L., D.S. performed CCR2 PET/CT experiments and Y.L. analyzed the PET/CT data. M.M. provided MC-21 antibody and technical expertise in experimental design. J.M., K.A., C.L., J.J., A.P. provided clinical expertise, assistance acquiring slides from human specimens for this study.

## Supplemental Material

Supplemental Methods

Figure S1-S18

Table S1

## Non-standard Abbreviations and Acronyms

ADCC: antibody-dependent cytotoxicity
AKT: RAC (Rho family)-alpha serine/threonine-protein kinase
BSA: Bovine serum albumin
Btg1: B-Cell Translocation Gene 1
Protein Cbr2: Carbonyl reductase 2
Ccl4: Chemokine (C-C motif) ligands 4
Ccl5: Chemokine (C-C motif) ligands 5
Ccl8: Chemokine (C-C motif) ligands 8
CCR2: C-C chemokine receptor type 2
CTLA4: cytotoxic T-lymphocyte-associated protein 4
CXCL10: Chemokine (C-X-C motif) ligand 10
CXCL16: Chemokine (C-X-C motif) ligand 16
CXCL9: Chemokine (C-X-C motif) ligand 9
CXCR3: CXC chemokine receptor 3
CXCR6: CXC chemokine receptor 3
DCM: dilated cardiomyopathy
DEG: Differentially expressed gene
FASL: Fas ligand
FCGR3A: Low affinity immunoglobulin gamma Fc region receptor III-A
Fcgr4: Fc receptor, IgG, low affinity IV
Folr2: Folate receptor beta
Gbp2b: Interferon-induced guanylate-binding protein 2b
GO: Gene ontology
HRP: Horseradish peroxidase
ICI: immune checkpoint inhibitor
ICM: ischemic cardiomyopathy
Icos: Inducible T-cell costimulator
IFN-γ: Interferon gamma
IrAEs: immune-related adverse events
KEGG: Kyoto Encyclopedia of Genes and Genomes
Klrd1: Killer cell lectin like receptor D1
Lag3: Lymphocyte Activating 3
Lars2: Probable leucyl-tRNA synthetase
Lgals3: Galectin-3
LM: lymphocytic myocarditis
LY-6C: lymphocyte antigen 6C
Lyve1: Lymphatic vessel endothelial hyaluronan receptor 1
MCP1: monocyte chemoattractant protein 1
MCP3: monocyte chemoattractant protein 3
MHC: major histocompatibility complex
MIF: macrophage migration inhibitory factor pathway
MYD88: Myeloid differentiation primary response 88
NK-cell: Natural Killer cell
Nlrp3: NLR family pyrin domain containing 3
Nrn1: Neuritin 1
PCA: Principal component analysis
PD1: Programmed cell death protein 1
PDL1: Programmed death-ligand 1
PET/CT: Positron emission tomography–computed tomography
Prf1: Perforin-1
SCENIC: Single-cell regulatory network inference and clustering
Stat1: Signal transducer and activator of transcription 1
TF: transcription factor
TNFα: Tumor necrosis factor α

## References

1. Wolchok JD, Kluger H, Callahan MK, Postow MA, Rizvi NA, Lesokhin AM, Segal NH, Ariyan CE, Gordon R-A and Reed K. Nivolumab plus ipilimumab in advanced melanoma. N Engl J Med. 2013;369:122–133.

2. Postow MA, Chesney J, Pavlick AC, Robert C, Grossmann K, McDermott D, Linette GP, Meyer N, Giguere JK and Agarwala SS. Nivolumab and ipilimumab versus ipilimumab in untreated melanoma. New England Journal of Medicine. 2015;372:2006–2017.

3. Motzer RJ, Tannir NM, McDermott DF, Frontera OA, Melichar B, Choueiri TK, Plimack ER, Barthélémy P, Porta C and George S. Nivolumab plus ipilimumab versus sunitinib in advanced renal-cell carcinoma. New England Journal of Medicine. 2018.

4. Heinzerling L, Ott PA, Hodi FS, Husain AN, Tajmir-Riahi A, Tawbi H, Pauschinger M, Gajewski TF, Lipson EJ and Luke JJ. Cardiotoxicity associated with CTLA4 and PD1 blocking immunotherapy. Journal for immunotherapy of cancer. 2016;4:1–11.

5. Varricchi G, Galdiero MR, Marone G, Criscuolo G, Triassi M, Bonaduce D, Marone G and Tocchetti CG. Cardiotoxicity of immune checkpoint inhibitors. ESMO open. 2017;2:e000247.

6. Johnson DB, Balko JM, Compton ML, Chalkias S, Gorham J, Xu Y, Hicks M, Puzanov I, Alexander MR and Bloomer TL. Fulminant myocarditis with combination immune checkpoint blockade. New England Journal of Medicine. 2016;375:1749–1755.

7. Sznol M, Ferrucci PF, Hogg D, Atkins MB, Wolter P, Guidoboni M, Lebbé C, Kirkwood JM, Schachter J and Daniels GA. Pooled analysis safety profile of nivolumab and ipilimumab combination therapy in patients with advanced melanoma. Journal of Clinical Oncology. 2017;35:3815–3822.

8. Wei SC, Meijers WC, Axelrod ML, Anang N-AA, Screever EM, Wescott EC, Johnson DB, Whitley E, Lehmann L and Courand P-Y. A genetic mouse model recapitulates immune checkpoint inhibitor–associated myocarditis and supports a mechanism-based therapeutic intervention. Cancer discovery. 2021;11:614–625.

9. Ji C, Roy MD, Golas J, Vitsky A, Ram S, Kumpf SW, Martin M, Barletta F, Meier WA and Hooper AT. Myocarditis in cynomolgus monkeys following treatment with immune checkpoint inhibitors. Clinical Cancer Research. 2019;25:4735–4748.

10. Wang J, Okazaki I-m, Yoshida T, Chikuma S, Kato Y, Nakaki F, Hiai H, Honjo T and Okazaki T. PD-1 deficiency results in the development of fatal myocarditis in MRL mice. International immunology. 2010;22:443–452.

11. Won T, Kalinoski HM, Wood MK, Hughes DM, Jaime CM, Delgado P, Talor MV, Lasrado N, Reddy J and Cihakova D. Cardiac myosin-specific autoimmune T cells contribute to immune-checkpoint-inhibitor-associated myocarditis. Cell Rep. 2022;41:111611.

12. Axelrod ML, Meijers WC, Screever EM, Qin J, Carroll MG, Sun X, Tannous E, Zhang Y, Sugiura A, Taylor BC, Hanna A, Zhang S, Amancherla K, Tai W, Wright JJ, Wei SC, Opalenik SR, Toren AL, Rathmell JC, Ferrell PB, Phillips EJ, Mallal S, Johnson DB, Allison JP, Moslehi JJ and Balko JM. T cells specific for alpha-myosin drive immunotherapy-related myocarditis. Nature. 2022;611:818–826.

13. Swirski FK and Nahrendorf M. Cardioimmunology: the immune system in cardiac homeostasis and disease. Nature Reviews Immunology. 2018;18:733–744.

14. Epelman S, Lavine KJ and Randolph GJ. Origin and functions of tissue macrophages. Immunity. 2014;41:21–35.

15. Bajpai G, Schneider C, Wong N, Bredemeyer A, Hulsmans M, Nahrendorf M, Epelman S, Kreisel D, Liu Y and Itoh A. The human heart contains distinct macrophage subsets with divergent origins and functions. Nature medicine. 2018;24:1234–1245.

16. Bajpai G, Bredemeyer A, Li W, Zaitsev K, Koenig AL, Lokshina I, Mohan J, Ivey B, Hsiao H-M and Weinheimer C. Tissue resident CCR2− and CCR2+ cardiac macrophages differentially orchestrate monocyte recruitment and fate specification following myocardial injury. Circulation research. 2019;124:263–278.

17. Lu H, Zong G, Zhou S, Jiang Y, Chen R, Su Z and Wu Y. Angiotensin II–C–C chemokine receptor2/5 axis-dependent monocyte/macrophage recruitment contributes to progression of experimental autoimmune myocarditis. Microbiology and immunology. 2017;61:539–546.

18. Leuschner F, Courties G, Dutta P, Mortensen LJ, Gorbatov R, Sena B, Novobrantseva TI, Borodovsky A, Fitzgerald K and Koteliansky V. Silencing of CCR2 in myocarditis. European heart journal. 2015;36:1478–1488.

19. Guerriero JL. Macrophages: Their Untold Story in T Cell Activation and Function. Int Rev Cell Mol Biol. 2019;342:73–93.

20. Archilla-Ortega A, Domuro C, Martin-Liberal J and Muñoz P. Blockade of novel immune checkpoints and new therapeutic combinations to boost antitumor immunity. Journal of Experimental & Clinical Cancer Research. 2022;41:1–24.

21. Finke D, Heckmann MB, Salatzki J, Riffel J, Herpel E, Heinzerling LM, Meder B, Völkers M, Müller OJ and Frey N. Comparative transcriptomics of immune checkpoint inhibitor myocarditis identifies guanylate binding protein 5 and 6 dysregulation. Cancers. 2021;13:2498.

22. Heo GS, Kopecky B, Sultan D, Ou M, Feng G, Bajpai G, Zhang X, Luehmann H, Detering L and Su Y. Molecular imaging visualizes recruitment of inflammatory monocytes and macrophages to the injured heart. Circulation research. 2019;124:881–890.

23. Moeller JB, Nielsen MJ, Reichhardt MP, Schlosser A, Sorensen GL, Nielsen O, Tornoe I, Gronlund J, Nielsen ME, Jorgensen JS, Jensen ON, Mollenhauer J, Moestrup SK and Holmskov U. CD163-L1 is an endocytic macrophage protein strongly regulated by mediators in the inflammatory response. J Immunol. 2012;188:2399–409.

24. Dick SA, Macklin JA, Nejat S, Momen A, Clemente-Casares X, Althagafi MG, Chen J, Kantores C, Hosseinzadeh S, Aronoff L, Wong A, Zaman R, Barbu I, Besla R, Lavine KJ, Razani B, Ginhoux F, Husain M, Cybulsky MI, Robbins CS and Epelman S. Self-renewing resident cardiac macrophages limit adverse remodeling following myocardial infarction. Nat Immunol. 2019;20:29–39.

25. Aibar S, González-Blas CB, Moerman T, Huynh-Thu VA, Imrichova H, Hulselmans G, Rambow F, Marine J-C, Geurts P and Aerts J. SCENIC: single-cell regulatory network inference and clustering. Nature methods. 2017;14:1083–1086.

26. Setty M, Kiseliovas V, Levine J, Gayoso A, Mazutis L and Pe’er D. Characterization of cell fate probabilities in single-cell data with Palantir. Nature biotechnology. 2019;37:451–460.

27. Mack M, Cihak J, Simonis C, Luckow B, Proudfoot AE, Plachy J, Bruhl H, Frink M, Anders HJ, Vielhauer V, Pfirstinger J, Stangassinger M and Schlondorff D. Expression and characterization of the chemokine receptors CCR2 and CCR5 in mice. J Immunol. 2001;166:4697–704.

28. Patel B, Bansal SS, Ismahil MA, Hamid T, Rokosh G, Mack M and Prabhu SD. CCR2(+) Monocyte-Derived Infiltrating Macrophages Are Required for Adverse Cardiac Remodeling During Pressure Overload. JACC Basic Transl Sci. 2018;3:230–244.

29. Rabin RL, Alston MA, Sircus JC, Knollmann-Ritschel B, Moratz C, Ngo D and Farber JM. CXCR3 is induced early on the pathway of CD4+ T cell differentiation and bridges central and peripheral functions. The Journal of Immunology. 2003;171:2812–2824.

30. Altara R, Mallat Z, Booz GW and Zouein FA. The CXCL10/CXCR3 axis and cardiac inflammation: implications for immunotherapy to treat infectious and noninfectious diseases of the heart. Journal of immunology research. 2016;2016.

31. Ni C, Ma P, Wang R, Lou X, Liu X, Qin Y, Xue R, Blasig I, Erben U and Qin Z. Doxorubicin-induced cardiotoxicity involves IFNγ-mediated metabolic reprogramming in cardiomyocytes. The Journal of Pathology. 2019;247:320–332.

32. Ma P, Qin Y, Cao H, Erben U, Ni C and Qin Z. Temporary blockade of interferon-γ ameliorates doxorubicin-induced cardiotoxicity without influencing the anti-tumor effect. Biomedicine & Pharmacotherapy. 2020;130:110587.

33. Weinmann SC and Pisetsky DS. Mechanisms of immune-related adverse events during the treatment of cancer with immune checkpoint inhibitors. Rheumatology. 2019;58:vii59-vii67.

34. Boughdad S, Latifyan S, Fenwick C, Bouchaab H, Suffiotti M, Moslehi JJ, Salem J-E, Schaefer N, Nicod-Lalonde M and Costes J. 68Ga-DOTATOC PET/CT to detect immune checkpoint inhibitor-related myocarditis. Journal for immunotherapy of cancer. 2021;9.

35. Luoma AM, Suo S, Williams HL, Sharova T, Sullivan K, Manos M, Bowling P, Hodi FS, Rahma O and Sullivan RJ. Molecular pathways of colon inflammation induced by cancer immunotherapy. Cell. 2020;182:655–671. e22.

36. Lavine KJ, Epelman S, Uchida K, Weber KJ, Nichols CG, Schilling JD, Ornitz DM, Randolph GJ and Mann DL. Distinct macrophage lineages contribute to disparate patterns of cardiac recovery and remodeling in the neonatal and adult heart. Proceedings of the National Academy of Sciences. 2014;111:16029–16034.

37. Epelman S, Lavine KJ, Beaudin AE, Sojka DK, Carrero JA, Calderon B, Brija T, Gautier EL, Ivanov S and Satpathy AT. Embryonic and adult-derived resident cardiac macrophages are maintained through distinct mechanisms at steady state and during inflammation. Immunity. 2014;40:91–104.

38. Salem JE, Bretagne M, Abbar B, Leonard-Louis S, Ederhy S, Redheuil A, Boussouar S, Nguyen LS, Procureur A, Stein F, Fenioux C, Devos P, Gougis P, Dres M, Demoule A, Psimaras D, Lenglet T, Maisonobe T, Pineton DECM, Hekimian G, Straus C, Gonzalez-Bermejo J, Klatzmann D, Rigolet A, Guillaume-Jugnot P, Champtiaux N, Benveniste O, Weiss N, Saheb S, Rouvier P, Plu I, Gandjbakhch E, Kerneis M, Hammoudi N, Zahr N, Llontop C, Morelot-Panzini C, Lehmann L, Qin J, Moslehi JJ, Rosenzwajg M, Similowski T and Allenbach Y. Abatacept/Ruxolitinib and Screening for Concomitant Respiratory Muscle Failure to Mitigate Fatality of Immune-Checkpoint Inhibitor Myocarditis. Cancer Discov. 2023.

39. Braster R, O’toole T and Van Egmond M. Myeloid cells as effector cells for monoclonal antibody therapy of cancer. Methods. 2014;65:28–37.

40. de Moel EC, Rozeman EA, Kapiteijn EH, Verdegaal EM, Grummels A, Bakker JA, Huizinga TW, Haanen JB, Toes RE and van der Woude D. Autoantibody development under treatment with immune-checkpoint inhibitors. Cancer Immunology Research. 2019;7:6–11.

41. Rikhi R, Karnuta J, Hussain M, Collier P, Funchain P, Tang WHW, Chan TA and Moudgil R. Immune checkpoint inhibitors mediated lymphocytic and giant cell myocarditis: uncovering etiological mechanisms. Frontiers in Cardiovascular Medicine. 2021;8.

42. Hokland P and Berg K. Interferon enhances the antibody-dependent cellular cytotoxicity (ADCC) of human polymorphonuclear leukocytes. The Journal of Immunology. 1981;127:1585–1588.

43. Tokunaga R, Zhang W, Naseem M, Puccini A, Berger MD, Soni S, McSkane M, Baba H and Lenz H-J. CXCL9, CXCL10, CXCL11/CXCR3 axis for immune activation–a target for novel cancer therapy. Cancer treatment reviews. 2018;63:40–47.

44. Zaidi MR. The interferon-gamma paradox in cancer. Journal of Interferon & Cytokine Research. 2019;39:30–38.

45. Salem J-E, Allenbach Y, Vozy A, Brechot N, Johnson DB, Moslehi JJ and Kerneis M. Abatacept for severe immune checkpoint inhibitor–associated myocarditis. New England Journal of Medicine. 2019;380:2377–2379.

46. Esfahani K, Buhlaiga N, Thébault P, Lapointe R, Johnson NA and Miller Jr WH. Alemtuzumab for immune-related myocarditis due to PD-1 therapy. New England Journal of Medicine. 2019;380:2375–2376.

47. Michel L, Helfrich I, Hendgen-Cotta UB, Mincu R-I, Korste S, Mrotzek SM, Spomer A, Odersky A, Rischpler C and Herrmann K. Targeting early stages of cardiotoxicity from anti-PD1 immune checkpoint inhibitor therapy. European heart journal. 2022;43:316–329.

