## Supplementary methods and figures for "Expansion of Disease Specific Cardiac Macrophages in Immune Checkpoint Inhibitor Myocarditis"

### Single-cell suspensions preparation and sorting

Heart tissue from *Ctla4*$^{+/-}$*Pdcd1*$^{-/-}$ and *Ctla4*$^{+/+}$*Pdcd1*$^{-/-}$ mice were minced with a razor blade and transferred into a 15 ml conical tube containing DMEM with Collagenase I (450 U/ml), DNAse I (60 U/ml), and Hyaluronidase (60 U/ml) and incubated at 37°C for 1 hour with agitation. Digestion was then stopped by addition of HBB buffer (2% FBS, 0.2% BSA in HBSS) and filtered through a 40 µm filter into 50 mL conical, transferred to a clean 15ml conical and centrifuged at 350 x G for 5 minutes at 4°C. Supernatant was then removed and pellet resuspended in 1 mL ACK lysing buffer (Gibco, A10492) and incubated at room temperature for 5 minutes followed by the addition of 9 mL DMEM. Suspension was then centrifuged at above conditions, followed by removal of supernatant, and resuspension in 5 mL FACS buffer (2% FBS, 2mM EDTA in calcium/magnesium free PBS). Centrifugation was repeated at above conditions, supernatant removed, and pellet resuspended in 300uL cell resuspension buffer (0.04% BSA in 1X PBS) and 1uL each of DRAQ5 (Thermo Scientific, 46162251) and DAPI (BD Biosciences, 564907) and allowed to incubate 5 minutes before sorting DRAQ5$^{+}$/DAPI$^{-}$ cells were collected in cell resuspension buffer. Collected cells were then recentrifuged according to above parameters and resuspended in cell resuspension buffer to a target concentration of 1000 cells/µL. Cells were counted on a hemocytometer and concentration adjusted as necessary.

### Single-cell RNA sequencing and analysis

All the codes used in the manuscript is deposited in GitHub. The Chromium Single Cell VDJ 5' Reagent V1.1 Kit from 10X Genomics was used to capture the transcriptome of individual cell. For each of the samples, 10,000 cells were loaded into a single well of a Chip G kit for GEM generation. Reverse transcription, barcoding, cDNA amplification, and purification were performed according to the Chromium 10x V1.1 protocol as the library preparation step. Each sample was sequenced on a NovaSeq 6000 S4 platform with a sequencing depth of approximately 20,000–30,000 reads per cell. Sequencing reads were aligned to the whole genome pre-mRNA reference generated from the mm10 transcriptome using the Cell Ranger V3 software (10X Genomics) according to the 10X Genomics instructions. The count matrices were pre-processed and analyzed using R package Seurat (v3.2.3). Cells with fewer than 1,000 or greater than 20,000 UMI counts, as well as cells with greater than 5 mitochondrial read percentage, were excluded from all subsequent analysis. 24556 cells remailed after quality control filtration and 97.7% (23988 cells) remained after doublet removal. For each sample, counts were transformed and normalized using SCTransform with default thresholds, and integration of the samples was performed on the filtered and normalized objects using Seurat v4, which identifies cells that have matching biological state across datasets based on canonical correlation analysis (CCA) and utilizes mutual nearest neighbors (MNN) to correct for batch effects. Principal component analysis (PCA) was performed on the integrated object using RunPCA. The number of significant PCs (ndims=20) was determined based on the ElbowPlot and DimHeatmap generated in Seurat. Non-linear dimensional reduction and visualization were performed using RunUMAP. Cells showing co-expression of multiple cell-type-specific genes were identified as doublets and removed from any downstream analysis. Differentially expressed genes were identified using the FindAllMarkers function on the normalized 'RNA' assay as recommended by Seurat, specifically returning only up-regulated genes with a $Log_2FC$ cutoff of 0.25 and a p-value of 0.05.

**Pathway enrichment analysis**

A list of output genes by Seurat different expression analysis with $Log_2FC>1$ and adjusted p-value<0.05 were used to identify enrichment in, myeloid and NK&T-cell clusters. Gene ontology (GO), Kyoto Encyclopedia of Genes and Genomes (KEGG) pathway analysis were performed using the ClusterProfiler package (version 2.4.3).

**Trajectory analysis**

Trajectory analysis was performed using the Python package 'Palantir'[1]. The count matrix and meta data sheet from the integrated object were exported as the input. Principal components and the diffusion components were calculated using the imported matrix as an estimate of the low dimensional phenotypic manifold of the data. Then, the Palantir algorithm was run with a specified start cell state (the presumed progenitor cell in the dataset). The algorithm then returned with the calculated terminal cells, entropy values, pseudotime values, and the probability of ending up in each of the terminal states for all cells.

**Cell-cell communication**

Cell to cell communication analysis was performed using the R package 'CellChat', which is a tool that is able to quantitatively infer and analyze intercellular communication networks from single-cell RNA-sequencing (scRNA-seq) data [2]. Basically, normalized expression matrix of macrophages and T-cells was inputted into a biological and statistical model. Based on a database of interactions among ligands, receptors and their cofactors that accurately represent known heteromeric molecular complexes, CellChat predicts major signaling inputs and outputs for cells and how those cells and signals coordinate for functions using network analysis and pattern recognition approaches.

**Transcription factor analysis**

SCENIC is a robust clustering method for the identification of stable cell states from scRNA-seq data based on the underlying gene regulatory networks (GRNs). It was performed as described[3, 4] (SCENIC 0.1.5, GENIE3 0.99.3 and AUCell 0.99.5) using the 10-thousand motifs database for RcisTarget (RcisTarget.mm9. motif Databases.10k). The input data was the size-factor normalized expression matrix in myeloid cells based on Seurat analysis filtered for Log2FC>0.5 and adjusted p-value<0.05. From those genes that passed the default filtering (rowSums > 5∗0.03∗760 and detected in at least 1% of the cells), only the protein coding genes were kept in the co-expression modules from GENIE3 and analyzed for motif enrichment with RcisTarget. AUCell score was used to visualize which cells are enriched for the gene set. TFs expression levels were visualized as colored projections onto UMAP. The primary results were further loaded as a matrix in R v4.0.1 for generation of heatmap and regulons network.

**Reference mapping and annotating query datasets**

We generated single cell RNA-seq dataset from $CD45^+$ cells isolated from wild type mouse hearts (n=6) and mapped the dataset onto the *Ctla4*$^{+/+}$*Pdcd1*$^{-/-}$ (n=4) and *Ctla4*$^{+/-}$ *Pdcd1*$^{-/-}$ (n=10) myeloid reference UMAP using the Azimuth workflow. The WT myeloid subset was selected as the query dataset. *Ctla4*$^{+/+}$*Pdcd1*$^{-/-}$ and *Ctla4*$^{+/-}$ *Pdcd1*$^{-/-}$ myeloid cells were selected as the reference. We computed mapping cell prediction scores of query cells to decipher the robustness of label transfer and mapping. Cell prediction scores reflect the confidence associated with each assigned annotation. Cells with high-confidence annotations (prediction scores > 0.75) reflect predictions that are supported by multiple consistent anchors.

**Immunofluorescence staining**

For human specimens, paraffin-embedded sections were dewaxed in xylene, rehydrated, endogenous peroxide activity quenched in 10% methanol and 3% hydrogen peroxide, processed for antigen retrieval by boiling in citrate buffer pH 6.0 for 15 mins, then blocked in 10% BSA containing 0.05% Tween-20, and stained with the following primary antibodies overnight at 4 °C: CD68 (KP1, Bio-Rad, 1:2,000), CD16α (SP189, Abcam, 1:200), CCR2 (7A7, Abcam,1:2000). The primary antibody was detected using Opal Polymer HRP Ms + Rb (PerkinElmer Opal Multicolor IHC system). For mouse samples, fresh frozen sections were endogenous peroxide activity quenched in 10% methanol and 3% hydrogen peroxide, blocked in 10% BSA containing 0.05% Tween-20, and stained with primary antibodies overnight at 4 °C: CD4 (EPR19514, Abcam, 1:1,000) , CD8 (D4W2Z, Cell Signaling Technology, 1:200) The primary antibody was detected using Opal Polymer HRP-Rb (PerkinElmer Opal Multicolor IHC system) .The PerkinElmer Opal Multicolor IHC system was utilized to visualize antibody staining per manufacturer protocol. For CD68 staining, fresh frozen sections were stained with primary antibody CD68 (FA-11, Biolegend, 1:100) and secondary goat anti rat-555 (A21434, Invitrogen,1:400). Immunofluorescence was visualized with Zeiss confocal microscopy and Zeiss Axio Scan Z1.

**Flow cytometry**

Saline perfused hearts were finely minced and digested with collagenase I (450 U/ml, Sigma CAS# 9001-12-1), hyaluronidase (60 U/ml, Sigma CAS# 37326-33-3) and DNase I (60 U/ml, Sigma CAS# 9003-98-9) for 1 hour at 37℃. Samples were then washed with Hanks' Balanced Salt Solution (Gibco, HBSS) that was supplemented with 2% Fetal Bovine Serum (Sigma, FBS) and 0.2% Bovine Serum Albumin (Sigma, BSA, CAS# 9048-46-8) and filtered through 40 μM cells trainers. Red blood cell lysis was performed with ACK lysis buffer (Thermo Fisher Scientific, cat# A1049201). Samples were washed with HBSS and resuspended in 100 μL of PBS with 2%

FBS and 2 mM EDTA (CORNING, product# 46-034-CI). Cells were stained with antibodies at 4℃ for 30 minutes. For IFN-γ staining, mice were injected with 250μg brefeldin (Biolegend, cat# 420601) i.p. 2 hours before sacrificing the mice and intracellular staining protocol followed the manufacturer protocol with eBioscience™ Foxp3 / Transcription Factor Fixation/Permeabilization Concentrate and Diluent kit (Thermo Fisher Scientific, cat# 00-5521-00). A complete list of antibodies is provided below. Samples were washed and resuspended. Flow cytometric analysis and sorting were performed on BD FACSMelody.

CD45-PerCP/Cy5.5, clone 30-F11 (Biolegend cat# 103131)

CD64-APC and PE, clone X54-5/7.1(Biolegend cat# 139305,139303)

CCR2-BV421, clone: SA203G11(Biolegend cat# 150605)

MHCII-APC/Cy7, clone M5/114.15.2 (Biolegend cat# 107627)

Ly6G-FITC, clone 1A8 (Biolegend cat# 127606)

Ly6C-BV510, clone HK1.4 (Biolegend cat# 128033)

IFN-γ-PE, clone XMG1.2 (Biolegend cat# 505807)

CD3e-FITC, clone145-2C11(ebioscience cat# 4338511)

CD19-BV510, clone 6D5(Biolegend cat# 115545)

CD49B-APC, clone DX5 (Biolegend cat# 108909)

CD16-2-PE/Cy7, clone 9E9(Biolegend cat# 149516)

CD45-BV510, clone 30-F11 (Biolegend cat# 103137)

CD4-PE/Cy7, clone GK1.5 (Biolegend cat# 100422)

CD8α-APC/Cy7, clone 53-6.7(Biolegend cat# 100714)

**Bone-marrow derived macrophages (BMDMs) generation and stimulation**

To generate bone-marrow derived macrophages (BMDMs), isolated bone marrow cells from mouse femurs and tibia were cultured for 6 days in DMEM supplemented with 10% FBS, 5% macrophage colony stimulating factor (M-CSF), 5% horse serum, 1% streptomycin and 1% sodium pyruvate. On day 6, the media was replaced with M-CSF free media for another 2 days. For IFN-γ stimulation, $1x10^6$ BMDMs were seeded on 6-well plates and cultured overnight to facilitate adherence. BMDMs were exposed to 50 ng/ml IFN-γ (R&D system cat# 485-MI-100/CF) with or without 25 μM Ruxolitinib (MCE, INCB18424). For Stat1 knockdown, BMDMs were transfected with 125 pM *Ctrl*siRNA (ThermoFisher, cat# AM4611) or *Stat1*siRNA (ThermoFisher, cat# AM16708) with Lipofectamine™ RNAiMAX Transfection Reagent (ThermoFisher, Cat# 13778030) and incubated for 48h for IFN-γ stimulation. For RT-PCR analysis, BMDMs were stimulated for 24h. For Western blot analysis, BMDMs were stimulated for 10min or 30min.

**Western blot**

To extract the total protein from mouse heart or BMDMs, collected heart tissue or cells were homogenized in RIPA lysis buffer containing protease inhibitor PMSF, and the total protein concentration were measured by BCA Protein Assay Kit (Thermo Fisher Scientific, cat#23225). 15μg protein samples were loaded. Primary antibodies: anti-α-Tubulin (Cell Signaling Technology, cat# 2144), STAT1(Cell Signaling Technology, cat#9172), p-STAT1 (Cell Signaling Technology, cat# 8242) and β-actin (Cell Signaling Technology, cat#3700) were incubated overnight at 4℃ and detected using peroxidase-conjugated goat anti-rabbit (Invitrogen, cat#31460) or goat anti-mouse (Invitrogen, cat#31430) as secondary antibody. Immuno-reactive bands were visualized with the Super Signal West Pico Chemiluminescent Substrate (Pierce). The intensity of target protein bands was quantified with Image J software (National Institutes of Health, USA) and normalized to Tubulin.

**RT-PCR**

RNA was extracted from myocardial tissue using the RNeasy RNA mini kit. RNA concentration was measured using a nanodrop spectrophotometer (Thermo Fisher Scientific). cDNA synthesis was performed using the HighCapacity RNA to cDNA synthesis kit (Applied Biosystems). Quantitative real time PCR reactions were prepared with sequence-specific primers with PowerUP™ Syber Green Master mix (Thermo Fisher Scientific) in a 20 μL volume. Real time PCR was performed using QuantStudio3 (Thermo Fisher Scientific). mRNA expression was normalized to 36B4. Primers were purchased from IDT. Primers sequences are provided below.

*Gbp2b*: ACAACTCAGCTAACTTTGTGGG; TGATACACAGGCGAGGCATATTA

*Cxcl9:* GTTCGAGGAACCCTAGTGATAAG; GTTTGAGGTCTTTGAGGGATTTG

*Cxcl10:* TCAGGCTCGTCAGTTCTAAGT; CCTTGGGAAGATGGTGGTTAAG

*Ccl8:* AGCTACGAGAGAATCAACAATATCC; CATGTACTCACTGACCCACTTC

*Fcgr4:* ATGTGGCAGCTACTACTACCA; ACCCACTTGGGGTCTAGGTTC

*Ifng:* GGCCATCAGCAACAACATAAG; GTTGACCTCAAACTTGGCAATAC

*Stat1:* CCACGCCTTTGGGAAGTATTA; GGTGGACTTCAGACACAGAAA

*36B4:* ATCCCTGACGCACCGCCGTGA; TGCATCTGCTTGGAGCCCACGTT

**RNAscope**

RNA was visualized using RNAScope Multiplex Fluorescent Reagent Kit v2 Assay, RNAScope 2.5 HD Detection Reagent-RED kits (Advanced Cell Diagnostics) using probes designed by Advanced Cell Diagnostics for *CXCL9*, *CXCL10*, *Cxcl9*, *Cxcl10, Ccr2, Cd68*. Paraffin section and fresh-frozen sections were used following indicated protocols. Chromogenic/brightfield images were acquired using a Zeiss Axio scan Z1 automated slide scanner. Image processing was performed using Zen Blue and Zen Black (Zeiss).

**CCR2$^+$ monocytes depletion**

To deplete CCR2$^+$ monocytes, mice were given 10 µg anti-CCR2 rat monoclonal antibody MC-21 or control antibody (clone LTF-2, BioXCell, cat# BP0090) i.p.[7] for five consecutive days starting at 28 days of age. For cardiac *Cxcl9*, *Cxcl10* in-situ mRNA signal detection and flow cytometry analysis, mouse heart was harvested 6 days after first MC-21 antibody treatment.

**CD8$^+$ T-cells depletion**

To deplete CD8$^+$ T-cells, mice were given 200 µg anti-CD8 rat monoclonal antibody (clone 2.43, BioXCell, cat# BP0061) or control antibody (clone LTF-2, BioXCell, cat# BP0090) i.p. every 3 days starting at 28 days of age. For cardiac *Cxcl9*, *Cxcl10* in-situ mRNA signal detection and flow cytometry analysis, mouse heart was harvested 6 days after first anti-CD8 treatment.

**IFN-γ signaling blockade**

Mice were injected i.p. with either 100 µL saline with 300 µg anti-IFN-γ (clone R46-A2; BioXCell, cat# BE0054) or rat IgG1 isotype control (clone HRPN, BioXCell, cat# BE0088), once a week, starting at 21 days of age. For survival time analysis, mortality was recorded daily during the whole experimental period. For cardiac *Cxcl9*, *Cxcl10* in-situ mRNA signal detection, mouse heart was harvested 23 days after first anti-IFN-γ treatment.

**Macrophage depletion**

To deplete $CD64^+$ macrophages, mice were given 300 μg anti-CSF1R rat monoclonal antibody (clone AFS98, BioXCell, cat# #BP0213) or control antibody (clone 2A3, BioXCell, cat# BP0089) i.p. every 2 days starting at 21 days of age. For survival time analysis, mortality was recorded daily during the whole experimental period. For cardiac *Cxcl9*, *Cxcl10* in-situ mRNA signal detection and flow cytometry analysis, mouse heart was harvested 23 days after first anti-CSF1R treatment.

**Randomization and Blinding**

For monocytes, $CD8^+$T-cells, macrophages depletion and IFN-γ blockade in vivo experiments, *Ctla4*$^{+/-}$*Pdcd1*$^{-/-}$mice were randomized (simple randomization) into two groups and treated with isotypes or MC-21, anti-CD8, anti-CSF1R, anti-IFN-γ, respectively. Blinding included injections, experiments, and analyses.

**CCR2 PET/CT imaging**

The CCR2 targeted PET tracer $^{68}$Ga-DOTA-ECL1i was synthesized and characterized as previously reported. Small animal PET/CT scans (40- to 60-min dynamic scan) was performed with Inveon PET/CT system (Siemens, Malvern, PA) on mice (6~8-week-old) after tail-vein injection. The PET images were corrected for attenuation, scatter, normalization, and camera dead time and co-registered with CT images. The PET images were reconstructed with the maximum a posteriori (MAP) algorithm and analyzed by Inveon Research Workplace. The heart uptake was calculated as the percent injected dose per gram (%ID/g) of tissue in three-dimensional regions of interest (ROIs) without the correction for partial volume effect[8, 9].

**References**

1. Setty M, Kiseliovas V, Levine J, Gayoso A, Mazutis L and Pe'er D. Characterization of cell fate probabilities in single-cell data with Palantir. *Nature biotechnology*. 2019;37:451-460.

2. Jin S, Guerrero-Juarez CF, Zhang L, Chang I, Ramos R, Kuan C-H, Myung P, Plikus MV and Nie Q. Inference and analysis of cell-cell communication using CellChat. *Nature communications*. 2021;12:1-20.

3. Aibar S, González-Blas CB, Moerman T, Huynh-Thu VA, Imrichova H, Hulselmans G, Rambow F, Marine J-C, Geurts P and Aerts J. SCENIC: single-cell regulatory network inference and clustering. *Nature methods*. 2017;14:1083-1086.

4. Aibar S, Gonzalez-Blas CB, Moerman T, Huynh-Thu VA, Imrichova H, Hulselmans G, Rambow F, Marine JC, Geurts P, Aerts J, van den Oord J, Atak ZK, Wouters J and Aerts S. SCENIC: single-cell regulatory network inference and clustering. *Nat Methods*. 2017;14:1083-1086.

5. Stuart T, Butler A, Hoffman P, Hafemeister C, Papalexi E, Mauck WM, 3rd, Hao Y, Stoeckius M, Smibert P and Satija R. Comprehensive Integration of Single-Cell Data. *Cell*. 2019;177:1888-1902 e21.

6. Wang YS, Chiu WT, Chang FP and Chen YL. Decline of chlorinated hydrocarbon insecticides residues in the tea-garden soils of Taiwan. *Proc Natl Sci Counc Repub China B*. 1988;12:9-13.

7. Bruhl H, Cihak J, Plachy J, Kunz-Schughart L, Niedermeier M, Denzel A, Rodriguez Gomez M, Talke Y, Luckow B, Stangassinger M and Mack M. Targeting of Gr-1+,CCR2+ monocytes in collagen-induced arthritis. *Arthritis Rheum*. 2007;56:2975-85.

8. Heo GS, Kopecky B, Sultan D, Ou M, Feng G, Bajpai G, Zhang X, Luehmann H, Detering L and Su Y. Molecular imaging visualizes recruitment of inflammatory monocytes and macrophages to the injured heart. *Circulation research*. 2019;124:881-890.

9. Heo GS, Bajpai G, Li W, Luehmann HP, Sultan DH, Dun H, Leuschner F, Brody SL, Gropler RJ and Kreisel D. Targeted PET imaging of chemokine receptor 2–positive monocytes and macrophages in the injured heart. *Journal of Nuclear Medicine*. 2021;62:111-114.

**Supplementary Figures and figure legends**

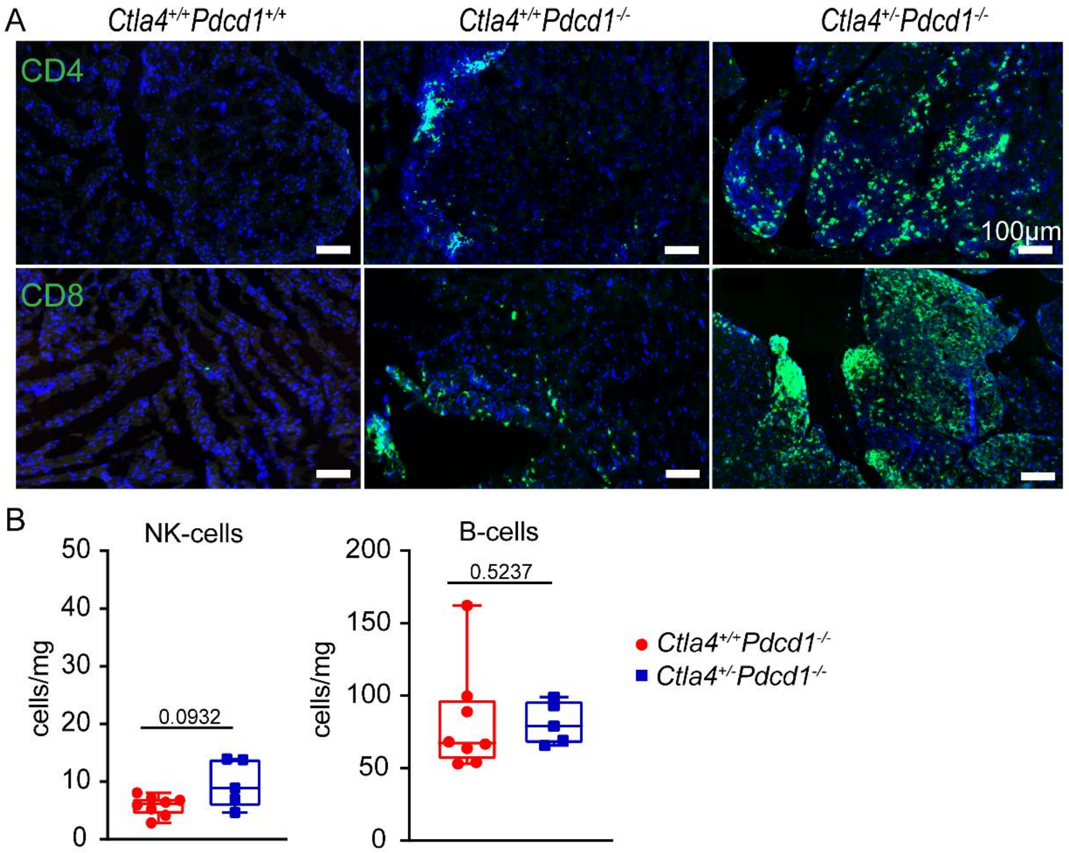


**Supplementary Fig 1. Accumulation of T-cells in *$Ctla4^{+/-}Pdcd1^{-/-}$* mouse hearts.** (A) Representative images of CD4 and CD8 fluorescent staining (green) heart sections. Scale bar, 100 µm. (B) Quantification of NK-cells, B-cells in the heart by flow cytometry. Data collected from two independent experiments. *$Ctla4^{+/+}Pdcd1^{-/-}$* (n=8), *$Ctla4^{+/-}Pdcd1^{-/-}$* (n=5), Mann-Whitney test, two-tailed.

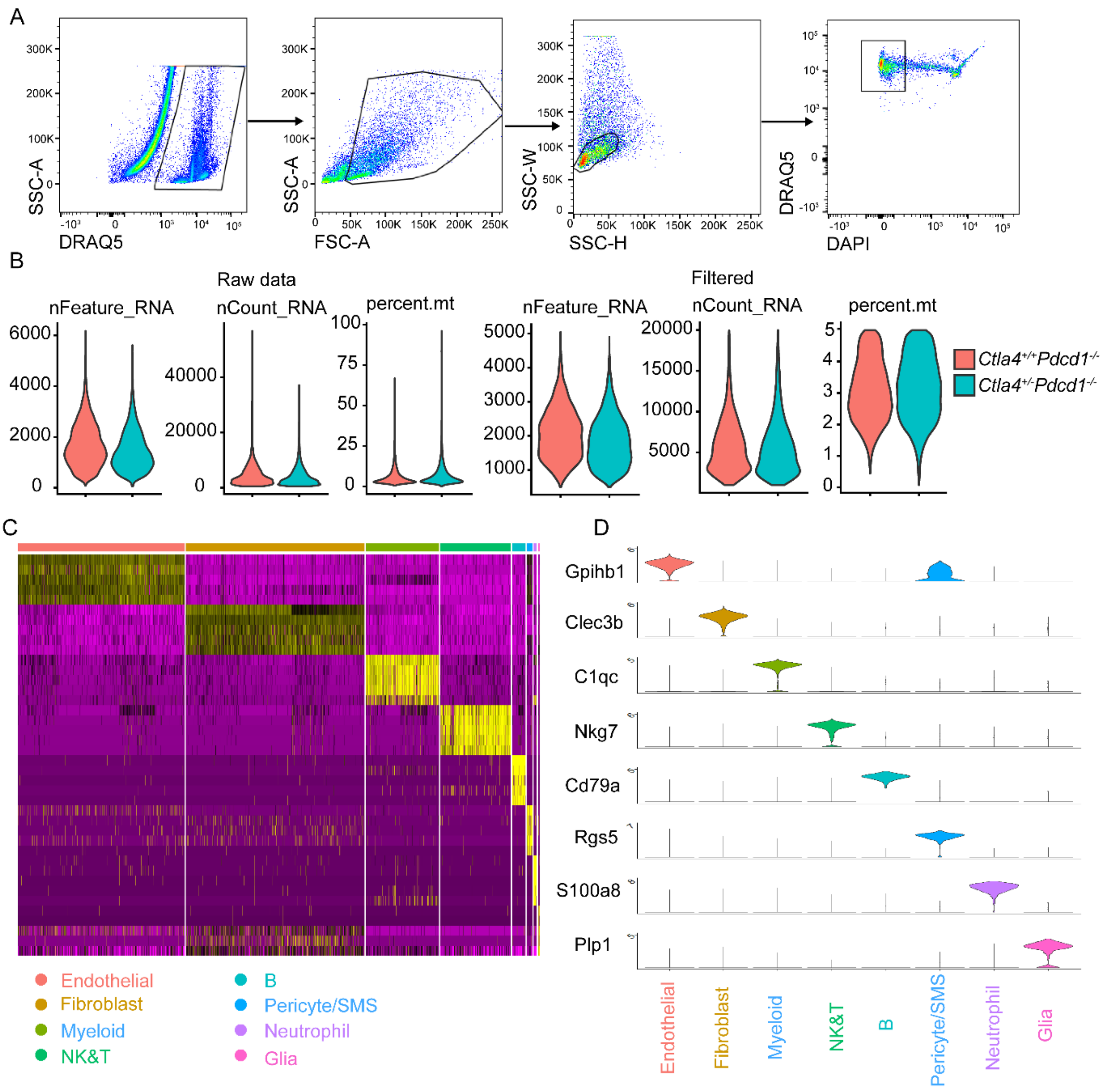


**Supplementary Fig 2. Quality control for single-cell RNA sequencing and gene expression signatures of major cell types.** (A) Flow cytometry cell sorting strategy for single cell RNA-sequencing. DRAQ5$^+$DAPI$^-$cells were sorted and subjected to single-cell RNA sequencing. (B) Violin plots show the number of features per cell (nFeature_RNA), number of unique molecular identifiers (UMIs) per cell (nCount_RNA), and percentage of mitochondrial reads per cell (percent.mt) before and after filtering. Cells with less than 1000 or more than 20,000 UMI counts, as well as cells with greater than 5% of mitochondrial reads were excluded from the study. (C) Heatmap shows relative expression profiles of distinct major cell types. Top bar and bottom legend indicate the populations. (D) Selected unique gene markers identified in each of the major cell types as shown in the violin plots.

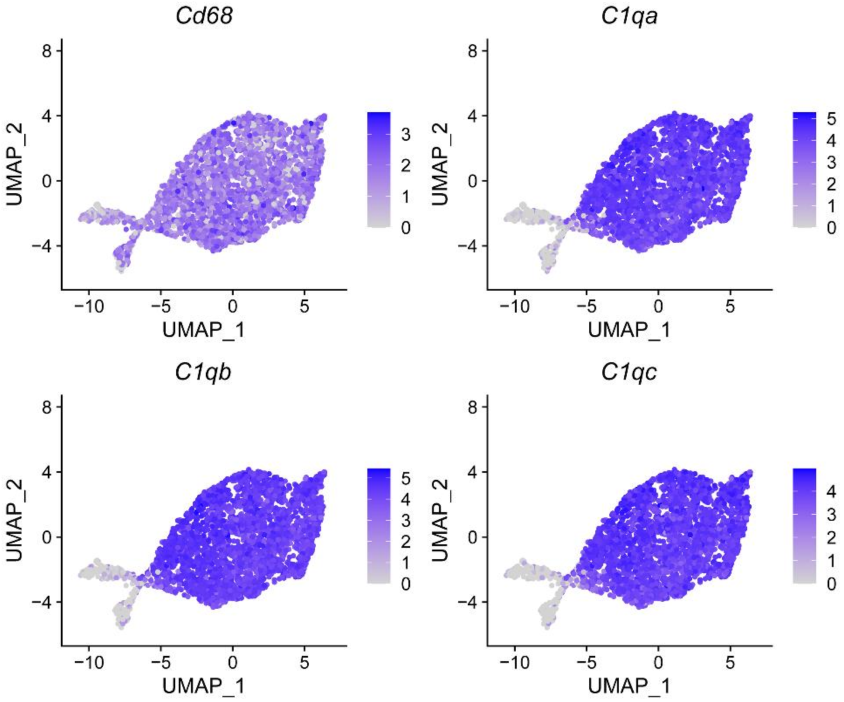


**Supplementary Fig 3. Macrophage marker gene expression.** Feature plots of macrophage marker genes (*Cd68, C1qa, C1qb, C1q*) in the myeloid cell subset.

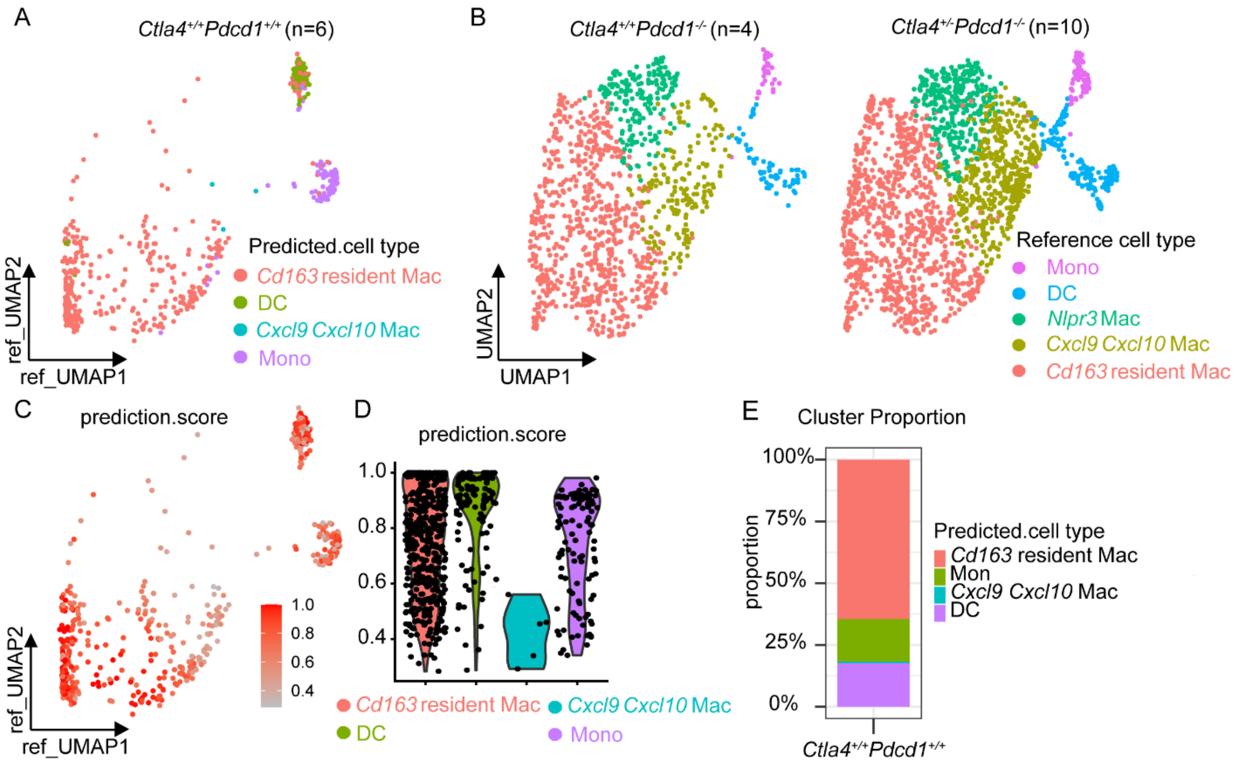


**Supplementary Fig 4. Reference mapping of myeloid subpopulations in query wide type *Ctla4*$^{+/+}$*Pdcd1*$^{+/+}$ dataset.** Reference mapping of query data (myeloid cells) from wide type mouse hearts (n=6) onto *Ctla4*$^{+/+}$*Pdcd1*$^{-/-}$ (n=4) and *Ctla4*$^{+/-}$ *Pdcd1*$^{-/-}$ (n=10) myeloid reference UMAP (B). (C) Feature plot and Violin plot (D) of prediction scores for various myeloid subpopulation in the query *Ctla4*$^{+/+}$*Pdcd1*$^{+/+}$ dataset. (E) The proportion of each myeloid subcluster in *Ctla4*$^{+/+}$*Pdcd1*$^{+/+}$ mice.

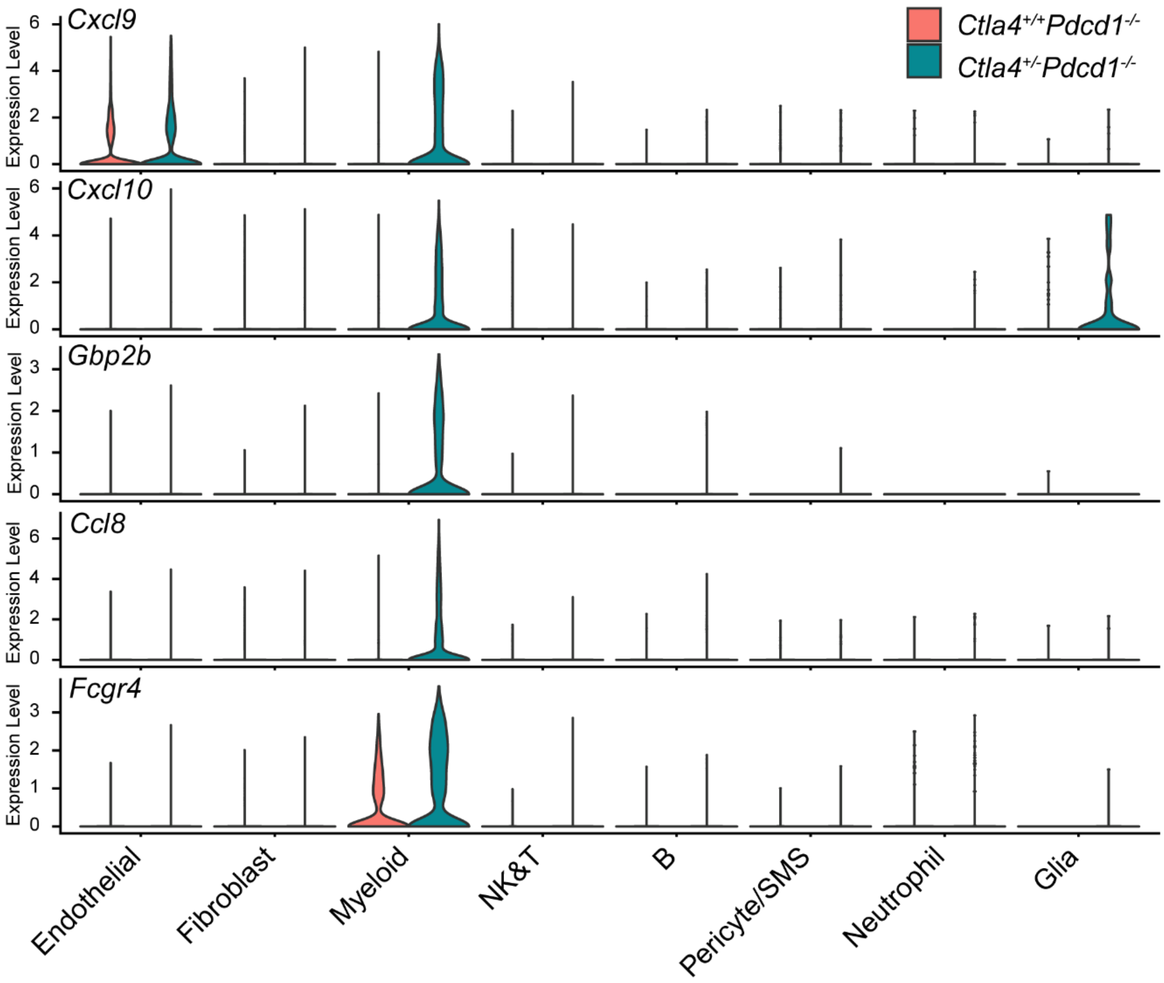


**Supplementary Fig 5. Enhanced expression of *Cxcl9, Cxcl10, Gbp2b, Ccl8, and Fcgr4* mRNA in *$Ctla4^{+/-}Pdcd1^{-/-}$* hearts within the myeloid cell subset.** Violin plot of *Cxcl9, Cxcl10, Gbp2b, Ccl8, and Fcgr4* mRNA expression in each of the major cell types.

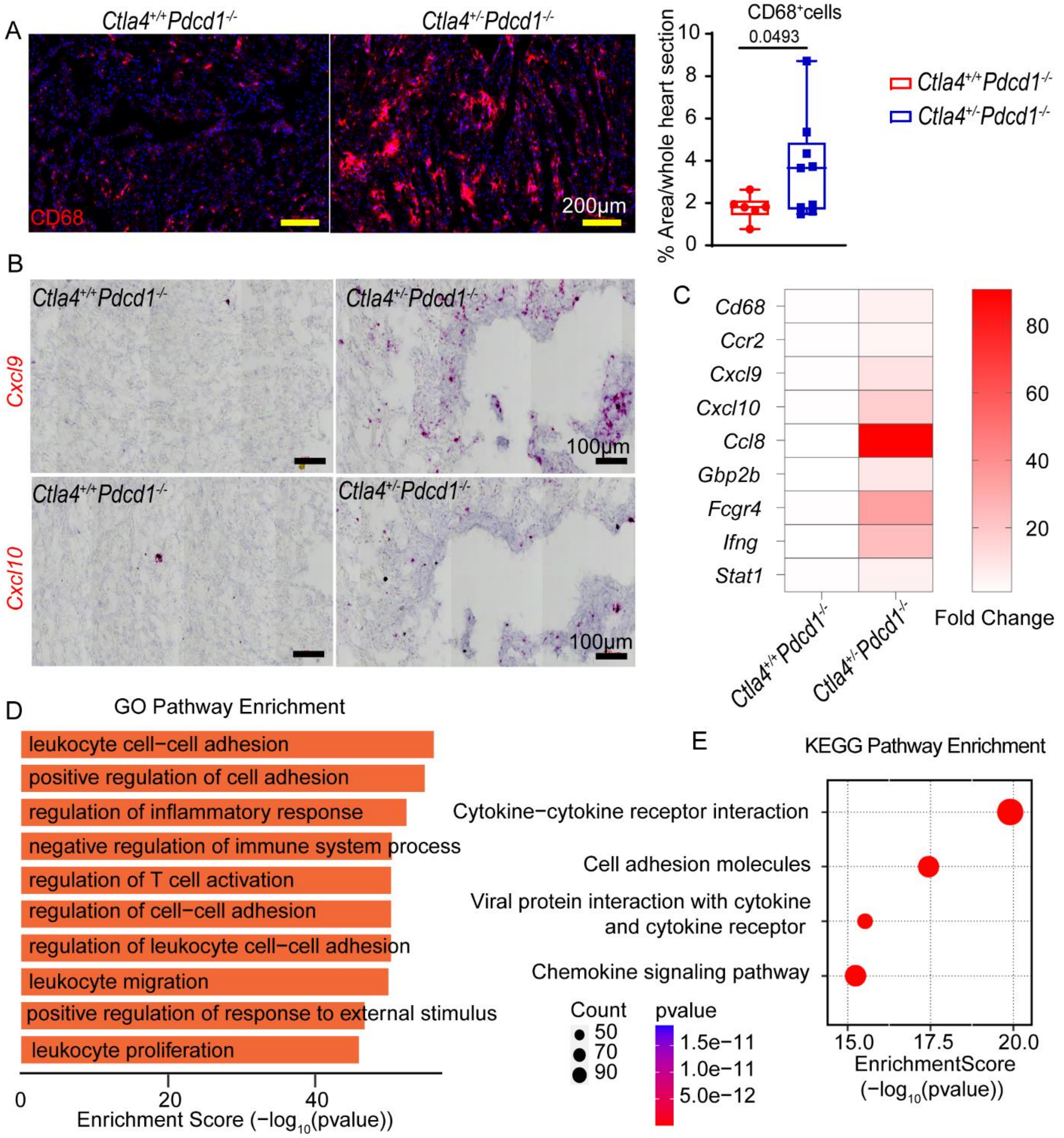


**Supplementary Fig 6. Accumulation of *Cxcl9 and Cxcl10* expressing cells in male *Ctla4*$^{+/-}$ *Pdcd1*$^{-/-}$ hearts.** (A) Representative images of CD68 immunofluorescent staining (red) in male *Ctla4*$^{+/+}$*Pdcd1*$^{-/-}$, and *Ctla4*$^{+/-}$*Pdcd1*$^{-/-}$ hearts. Quantification of CD68$^{+}$ cells. *Ctla4*$^{+/+}$*Pdcd1*$^{-/-}$ (n=6), *Ctla4*$^{+/-}$*Pdcd1*$^{-/-}$ (n=9), Welch's t test, two-tailed. Scale bar for CD68 staining images, 200 μm (B) Expression of *Cxcl9 and Cxcl10 mRNA was detected* in affected male mouse hearts via RNA in situ hybridization. *Representative images of Ctla4*$^{+/+}$*Pdcd1*$^{-/-}$ (n=5) and affected *Ctla4*$^{+/-}$*Pdcd1*$^{-/-}$ mice heart (2 of 7 mice) are shown. (C) Bulk cardiac expression of *Cd68/Ccr2/Cxcl9/Cxcl10/Ccl8/Gbp2b/Fcgr4/Ifng/Stat1* with RNA-seq data generated from male

*Ctla4*$^{+/+}$*Pdcd1*$^{-/-}$ (n=4) and affected *Ctla4*$^{+/-}$*Pdcd1*$^{-/-}$ (n=3) mice. Fold change was calculated based on RPKM value. (D-E) GO and KEGG pathway enrichment displaying the top enriched pathways in affected male *Ctla4*$^{+/-}$*Pdcd1*$^{-/-}$ *mice*. Genes used in the analysis were selected with $P < 0.05$ and $log_2FC > 1$. P-values were obtained by FDR corrected empirical Bayes moderated t-statistics in R package Limma. Pathway Enrichment P-values were calculated by hypergeometric distribution using R package ClusterProfiler.

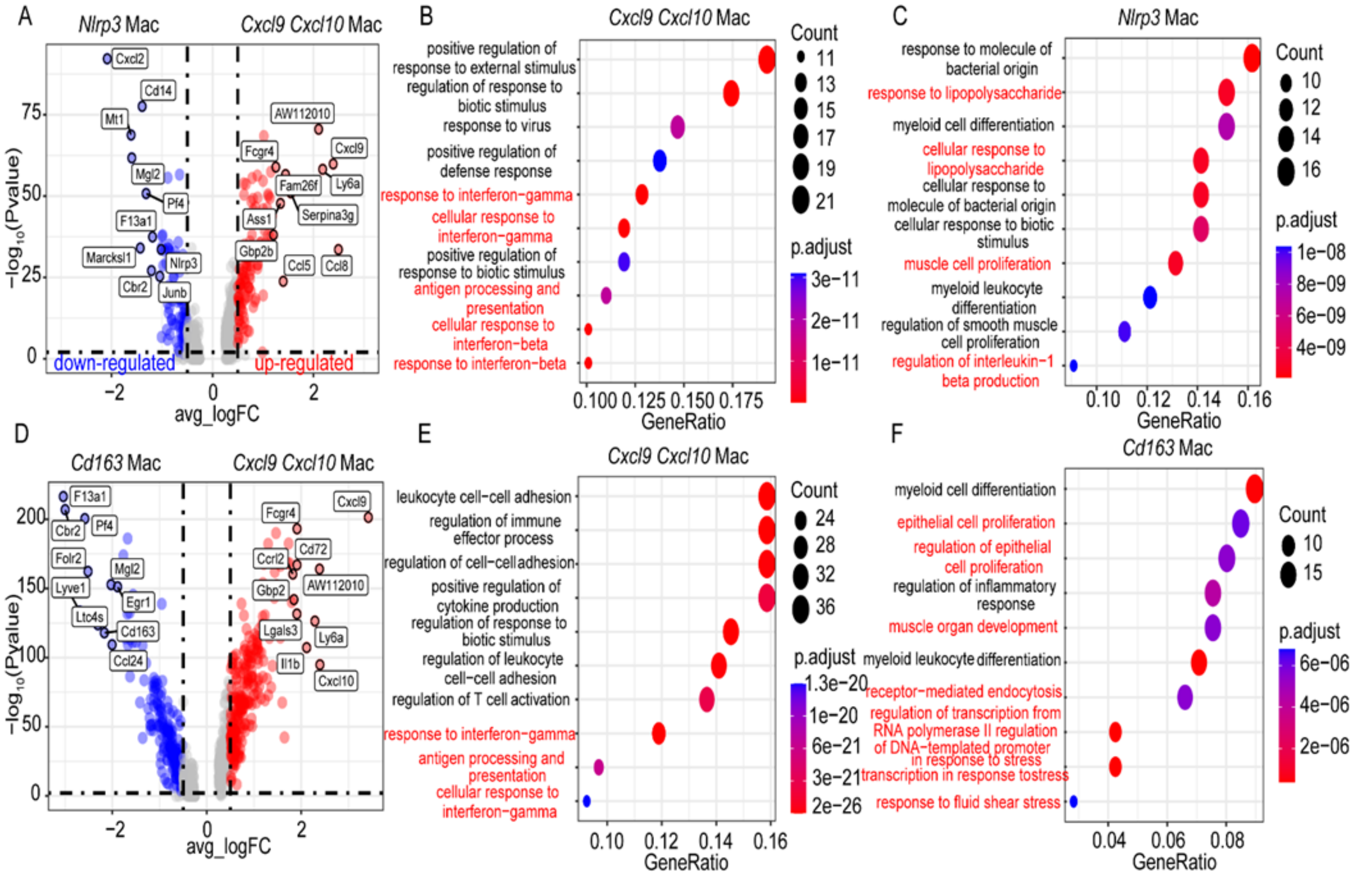


**Supplementary Fig 7. *Cxcl9*$^{+}$*Cxcl10*$^{+}$ macrophages exhibited distinct phenotypes compared with *Nlrp3*$^{+}$macrphages or *Cd163*$^{+}$ resident macrophages.** (A, D) Volcano plots showing differentially expressed genes (DEGs) between *Cxcl9*$^{+}$*Cxcl10*$^{+}$ macrophages vs. *Nlrp3*$^{+}$ macrophages and *Cxcl9*$^{+}$*Cxcl10*$^{+}$ macrophages vs. *Cd163*$^{+}$ resident macrophages. Gray: not significantly differentially expressed; blue: down regulated in *Cxcl9*$^{+}$*Cxcl10*$^{+}$ macrophages ($\log_2$ fold change>1, $p < 0.05$); red: upregulated in *Cxcl9*$^{+}$*Cxcl10*$^{+}$ macrophages ($\log_2$ fold change<-1, $p < 0.05$). obtained by Wilcoxon Rank Sum test using R package Seurat (v4). (B-C) GO pathway enrichment displaying the top enriched pathways in *Cxcl9*$^{+}$*Cxcl10*$^{+}$ macrophages vs. *Nlrp3*$^{+}$ macrophages. (D-E) GO pathway enrichment displaying the top enriched pathways in *Cxcl9*$^{+}$*Cxcl10*$^{+}$ macrophages vs. *Cd163*$^{+}$ resident macrophages. Genes used in the analysis selected from Seurat differential expression with $P < 0.05$ and $\log_2 FC > 1$. P value calculated using hypergeometric distribution and corrected for multiple comparisons.

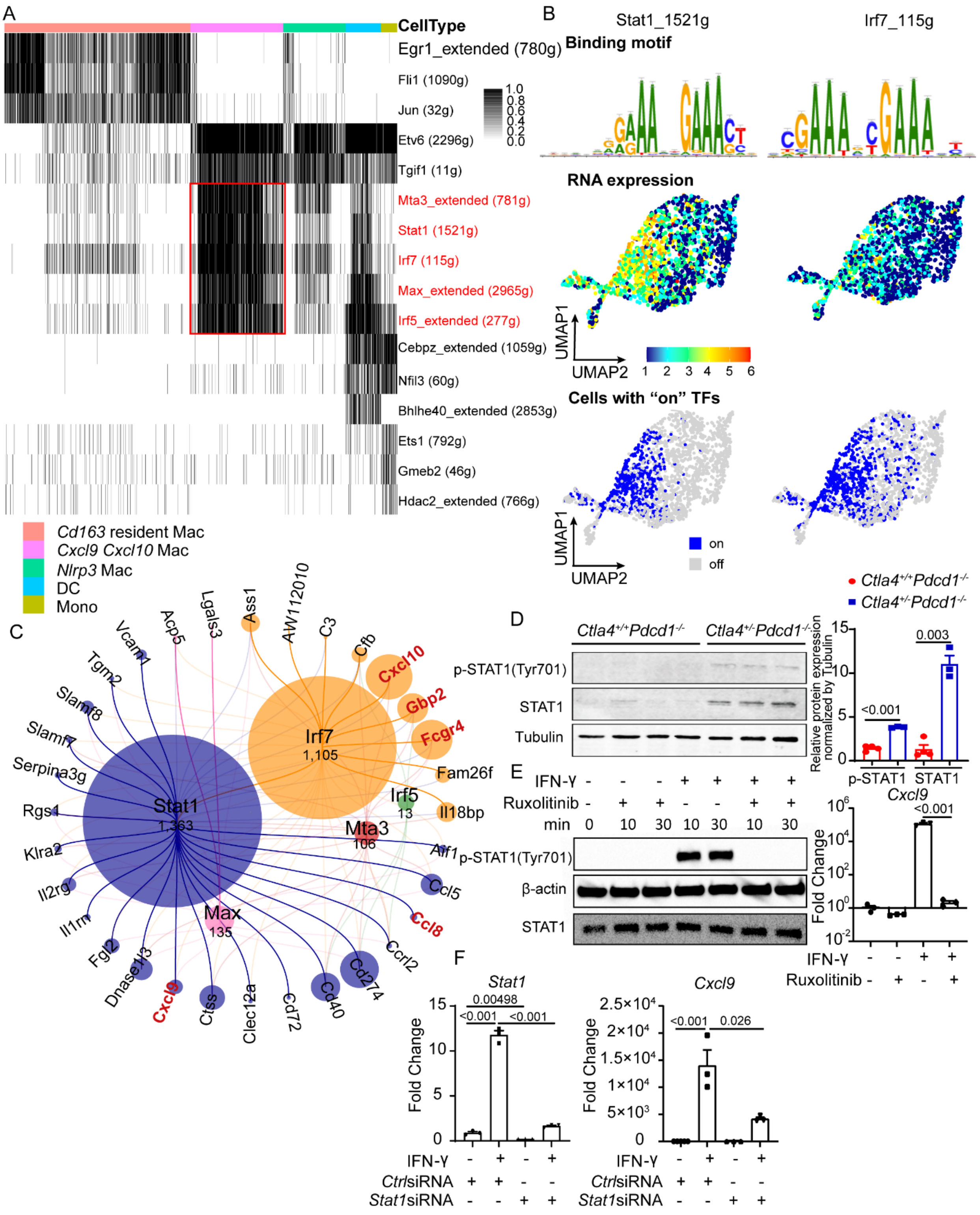


**Supplementary Fig 8. IFN-γ signaling induced the expansion of *Cxcl9*$^{+}$*Cxcl10*$^{+}$ macrophage via STAT1 activation.** (A) Binary heatmap generated from the activities of the transcription factor (TF) regulons for myeloid cells from *Ctla4*$^{+/-}$*Pdcd1*$^{-/-}$ mouse, with black blocks representing cells that are ‘on’ (activated TF regulon). The identity of cell type was assigned by established cluster information from the scRNA-seq data. Red boxes highlight the specific and activated TF regulons in *Cxcl9*$^{+}$*Cxcl10*$^{+}$ macrophages. Top bars show the cell type information. (B) Two regulons in *Cxcl9*$^{+}$*Cxcl10*$^{+}$ macrophages are highlighted: Stat1, Irf7. For each TF, Binding motif (upper) the RNA expression values of that TF (middle), and the cells passed the binary threshold set in the AUC histogram (lower) are shown. (C) A network generated with iRegulon using Stat1, Irf7, Irf5, Mta3, Max and the target genes as an input. The nodes, representing the TF regulons sized by the number of their motifs. The edges, representing the connections between each of the five TFs and their target genes, and shown as a line colored based on the TFs. Input target genes shown in the network are identified as marker genes for *Cxcl9*$^{+}$*Cxcl10*$^{+}$ macrophages. (D) Protein levels and phosphorylation states of STAT1 in *Ctla4*$^{+/+}$*Pdcd1*$^{-/-}$ and *Ctla4*$^{+/-}$*Pdcd1*$^{-/-}$ mouse heart was determined by western blot analysis. Welch's t-test or Unpaired t-test, two-tailed. (E)  BMDMs were exposed to IFN-γ with or without JAK1/2 inhibitor Ruxolitinib. (Left) Protein levels and phosphorylation states of STAT1 were indicated. (Right) Expression of *Cxcl9* was detected by RT-PCR. Welch's t-test or Unpaired t-test, two-tailed. (F) BMDMs were transfected with *Stat1* siRNA or Control siRNA and then stimulated with IFN-γ. Expression of *Stat1* and *Cxcl9* were detected by RT-PCR. Welch's t-test or Unpaired t-test, two-tailed.

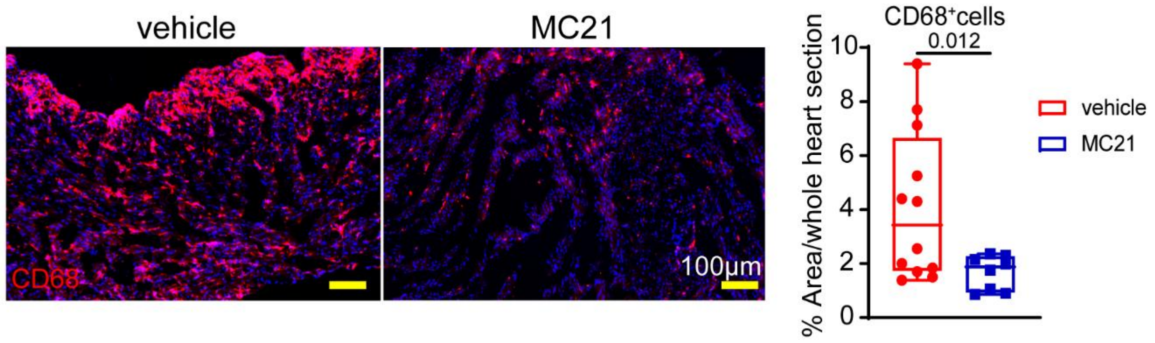


**Supplementary Fig 9. MC-21 treatment reduced the abundance of CD68$^{+}$cells in *Ctla4$^{+/-}$Pdcd1$^{-/-}$* mouse hearts.** Representative images of CD68 immunofluorescent staining (red) in vehicle or MC-21 treated *Ctla4$^{+/-}$Pdcd1$^{-/-}$* hearts. Quantification of CD68$^{+}$ cells in whole heart section. Vehicle (n=12), MC-21 (n=8), Welch's t test, two-tailed. Scale bar for CD68 staining images, 100 μm.

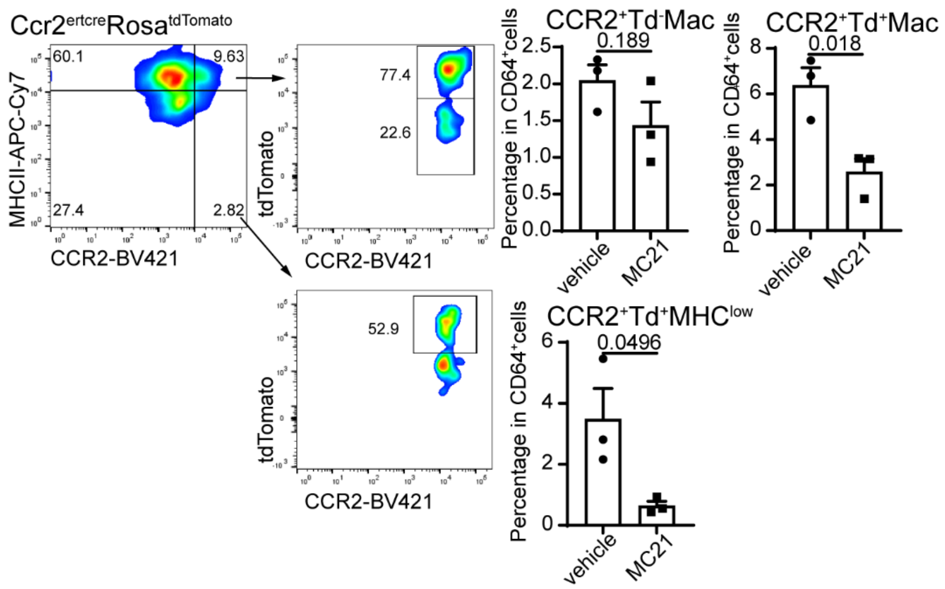


**Supplementary Fig 10. MC-21 antibody reduced CCR2$^{+}$macrophages by inhibition of monocytes recruitment not by direct effects on CCR2$^{+}$macropahges.** To label and track peripheral monocytes, *Ccr2$^{ertcre}$Rosa$^{tdTomato}$* mice were administrated tamoxifen (60mg/kg) by gavage every 3 days. Mice were then treated with *isotype or* MC-21 antibody. (Left) CCR2$^{+}$Td$^{-}$ macrophages represent CCR2$^{+}$ macrophages resident within the heart and CCR2$^{+}$Td$^{+}$ macrophages represent CCR2$^{+}$ macrophages that were recently derived from recruited monocytes. (Right) Quantification of the percentage of CCR2$^{+}$Td$^{-}$ macrophages, CCR2$^{+}$Td$^{+}$ macrophages and CCR2$^{+}$Td$^{+}$MHC$^{low}$ cells in CD64$^{+}$cells. Welch's t-test or Unpaired t-test, two-tailed.

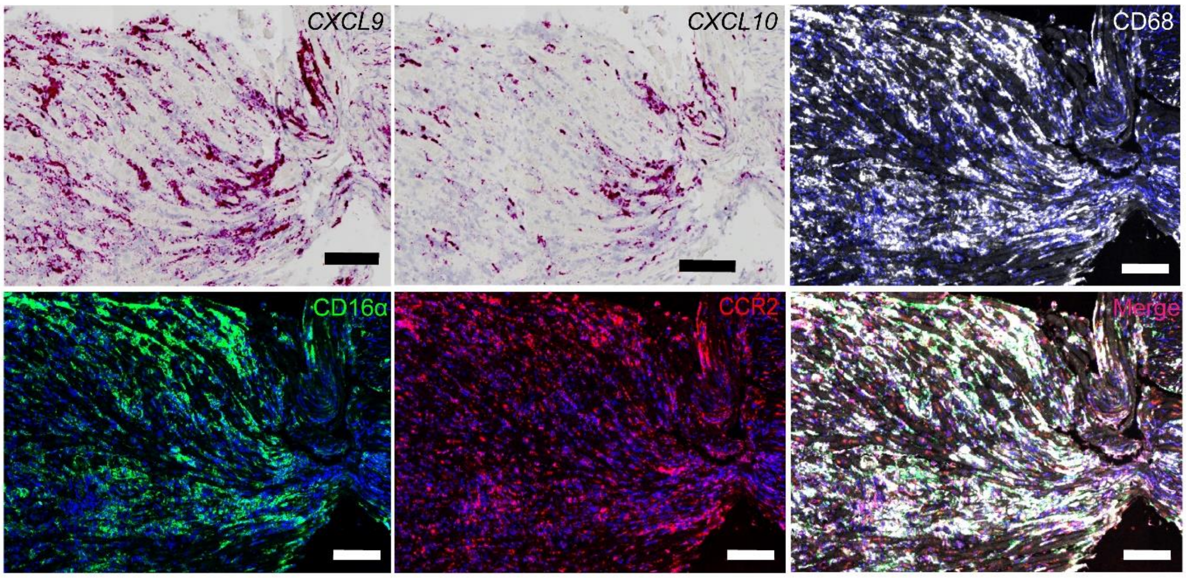


**Supplementary Fig 11. Co-localization of *CXCL9 and CXCL10 mRNA* and CD16α protein expression in macrophages within human ICI myocarditis specimens.** Expression of *CXCL9*, *CXCL10* via RNA in situ hybridization as well as immunofluorescence staining of CD16α (green), CCR2 (red), CD68 (white) and DAPI (blue) on consecutive sections from ICI myocarditis patient. scale bar, 100 µm.

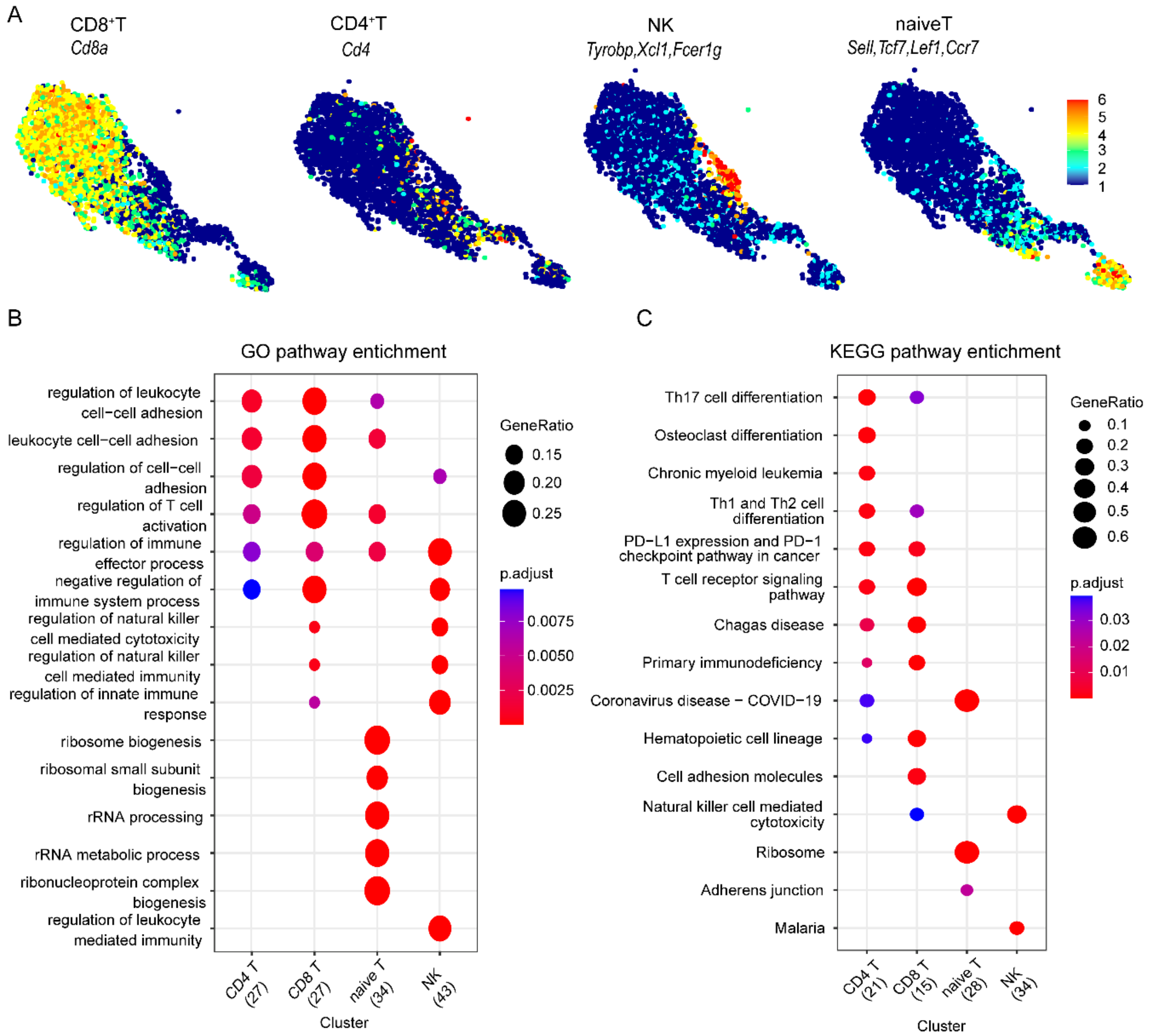


**Supplementary Fig 12. Marker genes and pathway analysis of NK cells and T cells** (A) Z score feature plots of marker genes for CD8 T-cells (*Cd8a*), CD4 T-cells (*Cd4*), naïve T-cells (*Sell, Tcf7, Lef1, Ccr7)*, and NK-cells (*Tyrobp, Xcl1, Fcer1g*). (B-C) GO and KEGG pathway enrichment displaying the top enriched pathways in each cluster. Genes used in the analysis selected from Seurat differential expression with P < 0.05 and $\log_2$FC > 1. P value calculated using hypergeometric distribution and corrected for multiple comparisons.

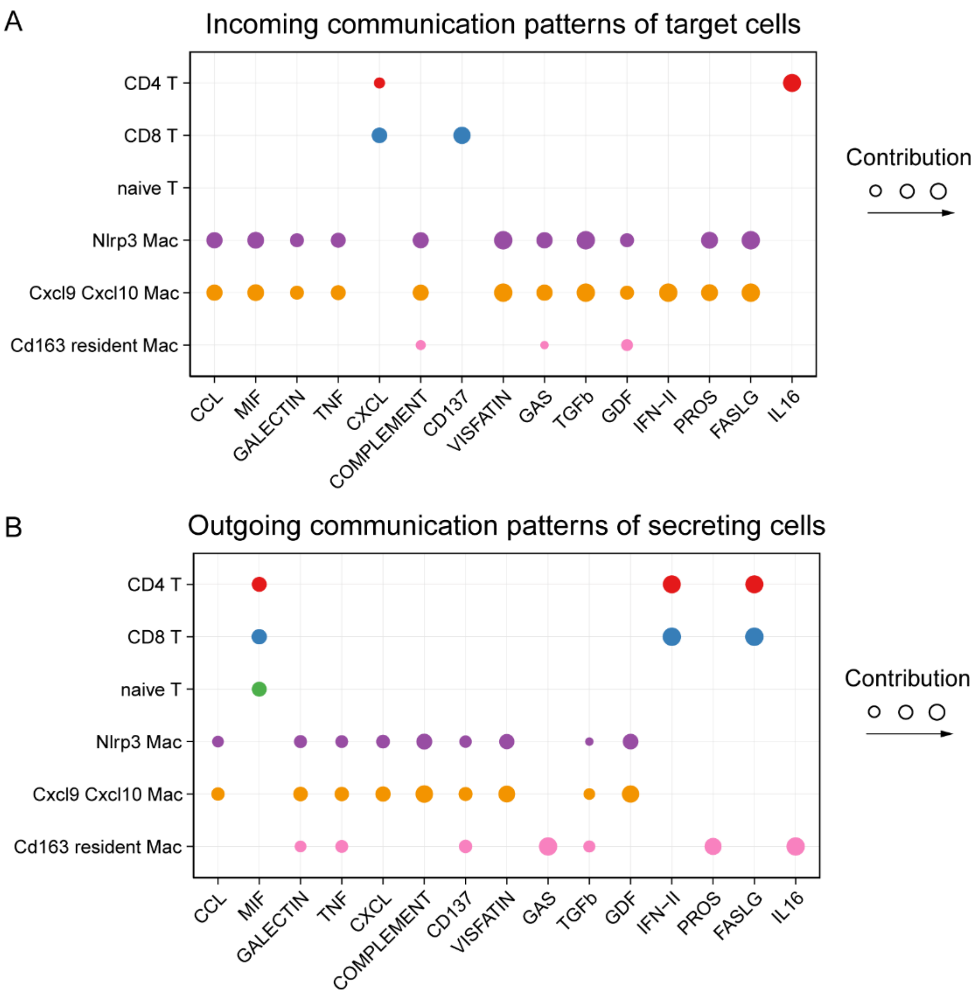


**Supplementary Fig 13. Comparator analysis of predicted communication between macrophages and T-cells.** Dot plots showing the comparison of incoming signaling patterns of target cells (A) and outgoing signaling patterns of secreting cells (B). The dot size is proportional to the contribution score computed from pattern recognition analysis. Higher contribution score implies the signaling pathway is enriched in the corresponding cell group.

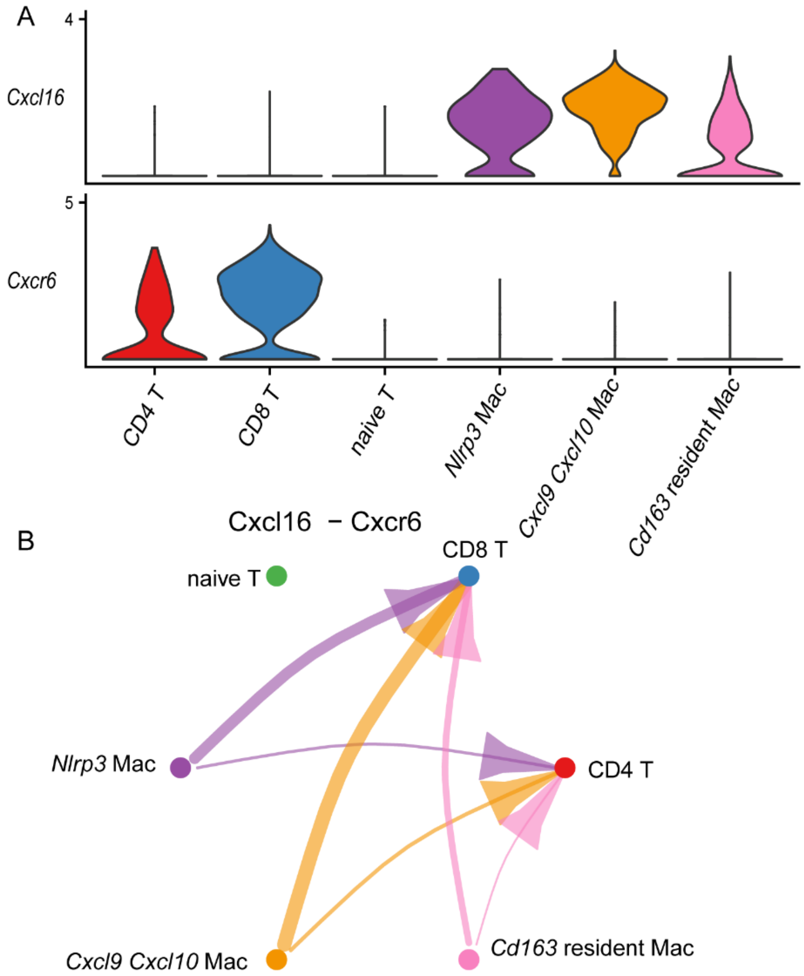


**Supplementary Fig 14. Cxcl16-Cxcr6 communication between macrophage and T-cell.** (A) Violin plot showing the expression distribution of signaling genes involved in the inferred Cxcl16-Cxcr6 interaction (B) Circle plot shows the inferred intercellular communication network for Cxcl16-Cxcr6 interaction.

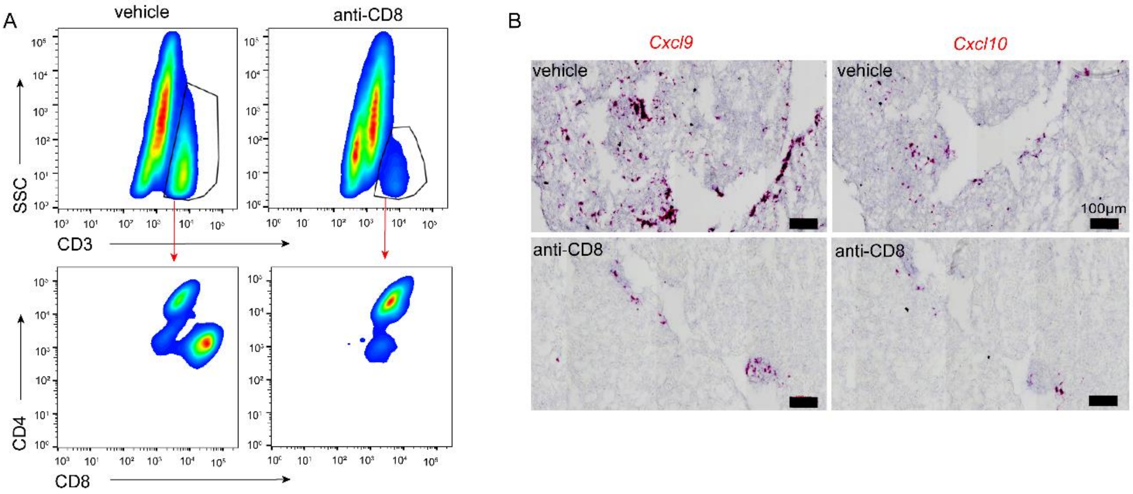


**Supplementary Fig 15. CD8$^+$ T-cell depletion suppresses the emergence of *Cxcl9*$^+$*Cxcl10*$^+$ macrophages.** (A) Cardiac CD8$^+$ T cells depletion was verified by flow cytometry. Displayed cells are CD45$^+$CD3$^+$. (B) Representative images of cardiac *Cxcl9* and *Cxcl10* mRNA expression 6 days after vehicle or anti-CD8 antibody treatment.

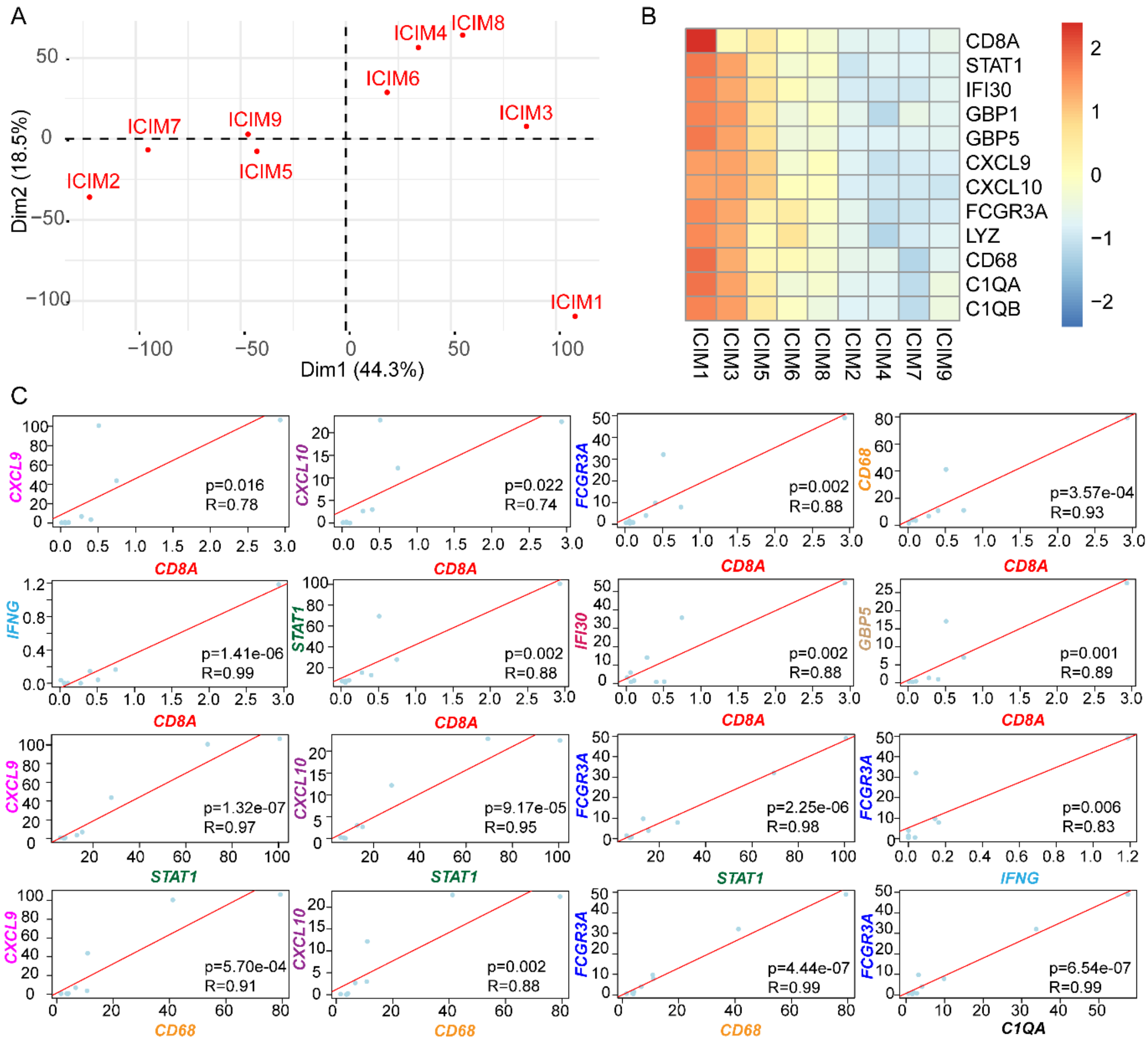


**Supplementary Fig 16. Correlation between the expression of *CD8A* and genes expressed in *Cxcl9*⁺*Cxcl10*⁺ macrophages in human ICI myocarditis biopsy samples.** (A) Principal Component Analysis (PCA) of transcriptome of human ICI myocarditis (ICIM) biopsy samples. Each dot represents a patient. (ICIM, 9 cases). (B) Heatmap of *CD8A, STAT1, IFI30, GBP1, GBP5, CXCL9, CXCL10, FCGR3A, LYZ, CD68, C1QA, and C1QB* mRNA expression. (C) Pearson correlation coefficient (r) and p-value (p) between the expression levels (TPM) of *CD8A* and *CXCL9, CXCL10, FCGR3A, CD68, IFNG, STAT1, IFI30, and GBP5*. Pearson correlation coefficient (r) and p-value (p) between the expression levels (TPM) of *STAT1 and CXCL9, CXCL10, and FCGR3A.* Pearson correlation coefficient (r) and p-value (p) between the expression levels (TPM) of *IFNG and FCGR3A.* Pearson correlation coefficient (r) and p-value (p) between the expression levels (TPM) *of CD68 and CXCL9, CXCL10, and FCGR3A.* Pearson correlation coefficient (r) and p-value (p) between the expression levels (TPM) of *C1QA and FCGR3A.*

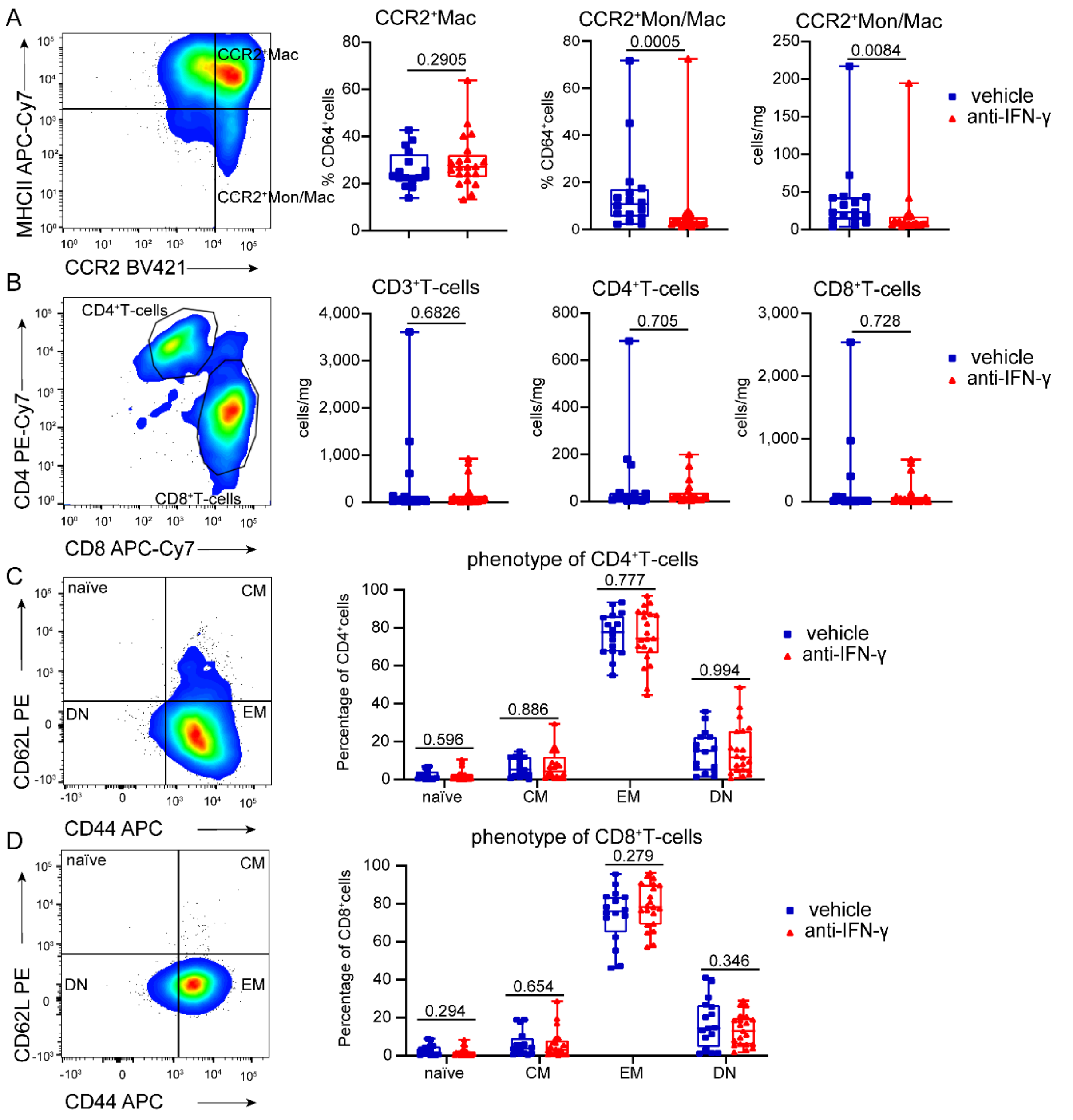


**Supplementary Fig 17. IFN-γ blockade results in reduced CCR2$^+$MHC$^{low}$ monocytes and macrophages.** (A) *Left*, Gating strategy of CD64$^+$ monocytes and macrophages. CCR2$^+$MHCII$^{high}$ cells correspond to CCR2$^+$ macrophages (CCR2$^+$ Mac) and CCR2$^+$MHC$^{low}$ cells are CCR2$^+$ monocytes and CCR2$^+$MHC$^{low}$ macrophages (CCR2$^+$ Mono/Mac). *Right*, Quantification of the percentage and absolute number of CCR2$^+$ Macs and CCR2$^+$Mon/Macs cells in the heart of *Ctla4*$^{+/-}$ *Pdcd1*$^{-/-}$ mice treated with vehicle (isotype control) or anti-IFN-γ antibody (R46A2) at 23 days after first antibody treatment. Percentages are relative to all $CD64^{+}$. Data collected from five independent experiments. vehicle (n=15); anti-IFN-γ (n=21), Mann-Whitney test, two-tailed. (B) *Left*, Gating strategy of $CD3^{+}$ T-cells. *Right*, Quantification of the absolute number of $CD3^{+}$ T-cells, $CD4^{+}$ T-cells, and $CD8^{+}$ T-cells in the heart of *Ctla4*$^{+/-}$ *Pdcd1*$^{-/-}$ mice treated with vehicle (isotype control) or anti-IFN-γ antibody (R46A2) at 23 days after first antibody treatment**.** Data collected from five independent experiments. vehicle (n=15); anti- IFN-γ (n=21), Mann-Whitney test, two-tailed. (C) *Left*, Gating strategy of $CD4^{+}$T-cells.$CD62L^{+}CD44^{-}$cells represent naïve T-cells; $CD62L^{+}CD44^{+}$cells represent central memory (CM) T-cells; $CD62L^{-}CD44^{-}$ cells represent double negative (DN) T-cells; $CD62L^{-}CD44^{+}$ cells represent effector memory (EM) T- cells. *Right*, quantification of the percentage of each $CD4^{+}$T-cell subset in the hearts of *Ctla4*$^{+/-}$*Pdcd1*$^{-/-}$ mice treated with vehicle (isotype control) or anti-IFN-γ antibody (R46A2) at 23 days after first antibody treatment. Data collected from five independent experiments. vehicle (n=15); anti-IFN-γ (n=21), Mann-Whitney test (naïve; CM; DN) or Welch's t test (EM), two-tailed. (D) *Left*, Gating strategy of $CD8^{+}$ T- cells. $CD62L^{+}CD44^{-}$ cells represent naïve T-cells; $CD62L^{+}CD44^{+}$ cells represent central memory (CM) T-cells; $CD62L^{-}CD44^{-}$ cells represent double negative (DN) T-cells; $CD62L^{-}CD44^{+}$ cells represent effector memory (EM) T-cells. *Right*, Quantification of the percentage of each $CD8^{+}$ T-cell subset in the hearts of *Ctla4*$^{+/-}$ *Pdcd1*$^{-/-}$ mice treated with vehicle (isotype control) or anti-IFN-γ antibody (R46A2) at 23 days after first antibody treatment**.** Data collected from five independent experiments. vehicle (n=15); anti-IFN-γ (n=21), Mann-Whitney test (naïve; CM) or Welch's t test (EM; DN), two-tailed.

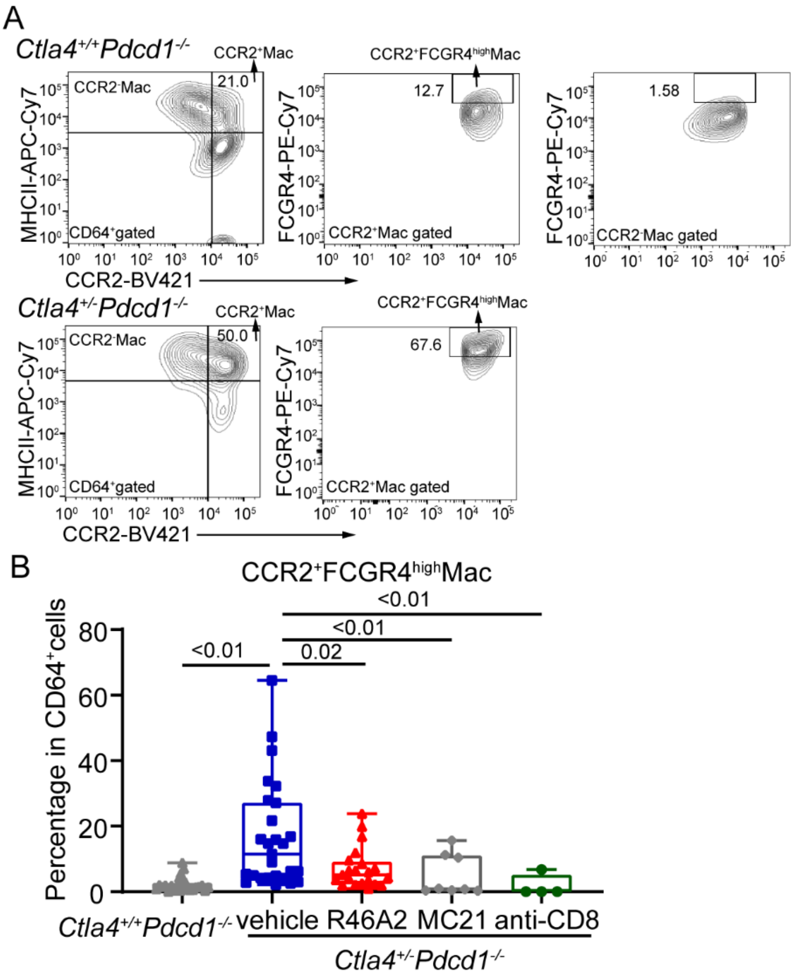


**Supplementary Fig 18. Anti-IFN-γ, MC-21 and anti-CD8 antibody treatment reduced the frequency of *Cxcl9*$^{+}$*Cxcl10*$^{+}$macrophages indicated by CCR2$^{+}$FCGR4$^{high}$macrophages using flow cytometry.** (A) Gating strategy of CCR2$^{+}$FCGR4$^{high}$macrophages (CCR2$^{+}$FCGR4$^{high}$Mac) in *Ctla4*$^{+/+}$*Pdcd1*$^{-/-}$ (Upper) and *Ctla4*$^{+/-}$*Pdcd1*$^{-/-}$ (Lower). CCR2$^{+}$FCGR4$^{high}$Mac correspond to activated *Cxcl9*$^{+}$*Cxcl10*$^{+}$macrophages. We use the expression level of FCGR4 in CCR2$^{-}$macrophages as baseline. (B) Quantification of the percentage of CCR2$^{+}$FCGR4$^{high}$Mac in the heart of *Ctla4*$^{+/+}$*Pdcd1*$^{-/-}$, *Ctla4*$^{+/-}$*Pdcd1*$^{-/-}$ mice treated with vehicle (isotype controls) or anti-IFN-γ (R46A2) or MC-21 or anti-CD8 antibody. Percentages are relative to all CD64$^{+}$cells. Vehicle (n=27); anti-IFN-γ (R46A2) (n=21), MC-21 (n=8), anti-CD8 (n=4), Mann-Whitney test, two-tailed.
